# Pharmacological Induction of mesenchymal-epithelial transition chemosensitizes breast cancer cells and prevents metastatic progression

**DOI:** 10.1101/2023.04.19.537586

**Authors:** Meisam Bagheri, Gadisti Aisha Mohamed, Mohammed Ashick, Mohamed Saleem, Nevena B. Ognjenovic, Hanxu Lu, Fred W. Kolling, Owen M. Wilkins, Subhadeep Das, Ian S. La Croix, Shivashankar H. Nagaraj, Kristen E. Muller, Scott A. Gerber, Todd W. Miller, Diwakar R. Pattabiraman

## Abstract

The epithelial-mesenchymal transition (EMT) is a developmental program co-opted by tumor cells that aids the initiation of the metastatic cascade. Tumor cells that undergo EMT are relatively chemoresistant, and there are currently no therapeutic avenues specifically targeting cells that have acquired mesenchymal traits. We show that treatment of mesenchymal-like triple-negative breast cancer (TNBC) cells with the microtubule-destabilizing chemotherapeutic eribulin, which is FDA-approved for the treatment of advanced breast cancer, leads to a mesenchymal-epithelial transition (MET). This MET is accompanied by loss of metastatic propensity and sensitization to subsequent treatment with other FDA-approved chemotherapeutics. We uncover a novel epigenetic mechanism of action that supports eribulin pretreatment as a path to MET induction that curtails metastatic progression and the evolution of therapy resistance.

**Summary:** While the advent of targeted therapy has led to vast improvements in outcomes for certain types of breast cancer, a mainstay for triple-negative breast cancer (TNBC) remains cytotoxic chemotherapy. A major clinical hurdle in successfully managing this disease is the eventual development of therapeutic resistance and disease relapse in more aggressive forms. Our data reveal that epigenetic modulation of EMT state using the FDA-approved therapeutic eribulin curtails metastatic propensity of breast tumors and, when administered in the treatment-naïve setting, sensitizes to subsequent treatment with other chemotherapeutics.

## Introduction

The epithelial-mesenchymal transition (EMT) is a cellular program that imparts plasticity to epithelial cells, enabling them to acquire traits including motility and invasion ^1^. While being an integral step in developmental processes such as gastrulation and neural crest migration as well as normal processes such as wound healing, the role of EMT in tumor progression has been studied in detail. In addition to contributing to the local invasion of primary tumor cells and enabling intravasation into blood/lymphatic vessels, EMT plays a crucial role in tumor cell extravasation and chemoresistance ^1,2^. Despite our deep understanding of EMT and the signaling pathways and transcriptional networks that regulate this plasticity in diverse cellular contexts, the impact of EMT research on the therapeutic targeting of carcinomas has been minimal. While there are FDA-approved therapies that can modulate EMT, there are no approved drugs administered for their ability to inhibit or reverse EMT. Indeed, there is a gap between (a) our biological understanding of EMT, and (b) translation of that knowledge to improve clinical outcomes.

Eribulin is a microtubule dynamics inhibitor administered to patients with advanced/metastatic breast cancer in the ≥3^rd^-line setting ^3^. These patients have often received prior chemotherapy that included an anthracycline and a taxane. The proposed primary mechanism of action or eribulin is the inhibition of tubulin addition to the plus end of microtubules, thereby inducing mitotic blockade ^4^. Through a seemingly unconnected molecular mechanism, eribulin is also known to induce mesenchymal-to-epithelial transition (MET) in some models ^5^. Although clues on the induction of MET have emerged from studies showing that eribulin can cause membrane localization of E-cadherin and inhibit TGFb-induced expression of Snail ^6,7^, the mechanism through which this purported microtubule dynamics inhibitor induces a global change in cell state remains unclear.

Using a combination of TNBC cells, genetically engineered mouse models, and patient-derived xenografts (PDX), we uncovered a novel mechanism through which Zeb1 interacts with members of the SWI/SNF family of chromatin remodelers to maintain the mesenchymal state. This interaction, when disrupted using pharmacologically achievable concentrations of eribulin, induces MET and alters the differentiation state of tumors. This MET is accompanied by a reduction in tumor-initiation and metastatic propensity of tumor cells, while also rendering them more susceptible to subsequent chemotherapy. Using a combination of lentiviral barcoding and single-cell analysis, we uncover that disrupting the Zeb1-SWI/SNF interaction using eribulin reprograms the chromatin and transcriptional profile of tumor cells, shifting their evolutionary trajectory as they progress through the bottleneck of treatment. This work sheds new light on our understanding of how cells utilize optimal chromatin remodeling machinery to maintain a more mesenchymal state that imparts aggressive traits while unveiling a novel mechanism of action for eribulin to modulate cancer cell state, paving the way for optimal clinical use exploiting its multi-faceted anti-tumor properties.

## Results

### Therapeutic induction of MET in breast cancer cells

Given the previously demonstrated role of eribulin in the modulation of EMT state ^5^, we sought to compare its effects with those of other chemotherapeutics within the same class of microtubule dynamics inhibitors. Paclitaxel is used in neoadjuvant chemotherapy for breast cancer, and vinorelbine, like eribulin, is administered to treat advanced breast cancers. All 3 drugs inhibited the growth of PB3 cancer cells derived from a MMTV-PyMT transgene-driven murine mammary (Fig. 1A), which reside in a quasi-mesenchymal state ^8^. To mimic clinical treatment regimens in the development of drug-resistant cell lines, chemotherapeutics were applied intermittently to PB3 cells for 3 cycles, allowing for intervening drug holidays for cells to recover (Fig. 1B).Treatment with eribulin at its IC_50_, but not with vinorelbine or paclitaxel at their IC_50_, led to the emergence of an eribulin-resistant cell population (ERI-R) exhibiting upregulated levels of epithelial cell markers [i.e., epithelial cell adhesion marker (EpCAM), E-cadherin], downregulated levels of mesenchymal markers (Vimentin, Zeb1), and an epithelial cobblestone-like morphology compared to PB3 controls, vinorelbine-resistant derivatives (VIN-R), and paclitaxel-resistant derivatives (PAC-R) (Fig. 1C–F). The altered phenotype of ERI-R cells was accompanied by decreased migratory and invasive potentials compared to control PB3 cells (Fig. 1G, H and Suppl. Fig.S1A). Importantly, chemotherapeutics were removed from drug-resistant cells 4 weeks prior to seeding for assays, indicating that these phenotypes were stable and not due to the continued presence of drugs. Similar effects of eribulin treatment were observed in human triple-negative breast cancer (TNBC) cells. ERI-R derivatives of MDA-MB-231 and SUM159PT cells showed increased E-cadherin and decreased VIM and ZEB1 (Suppl. Fig. S1B-I).

**Figure 1:**
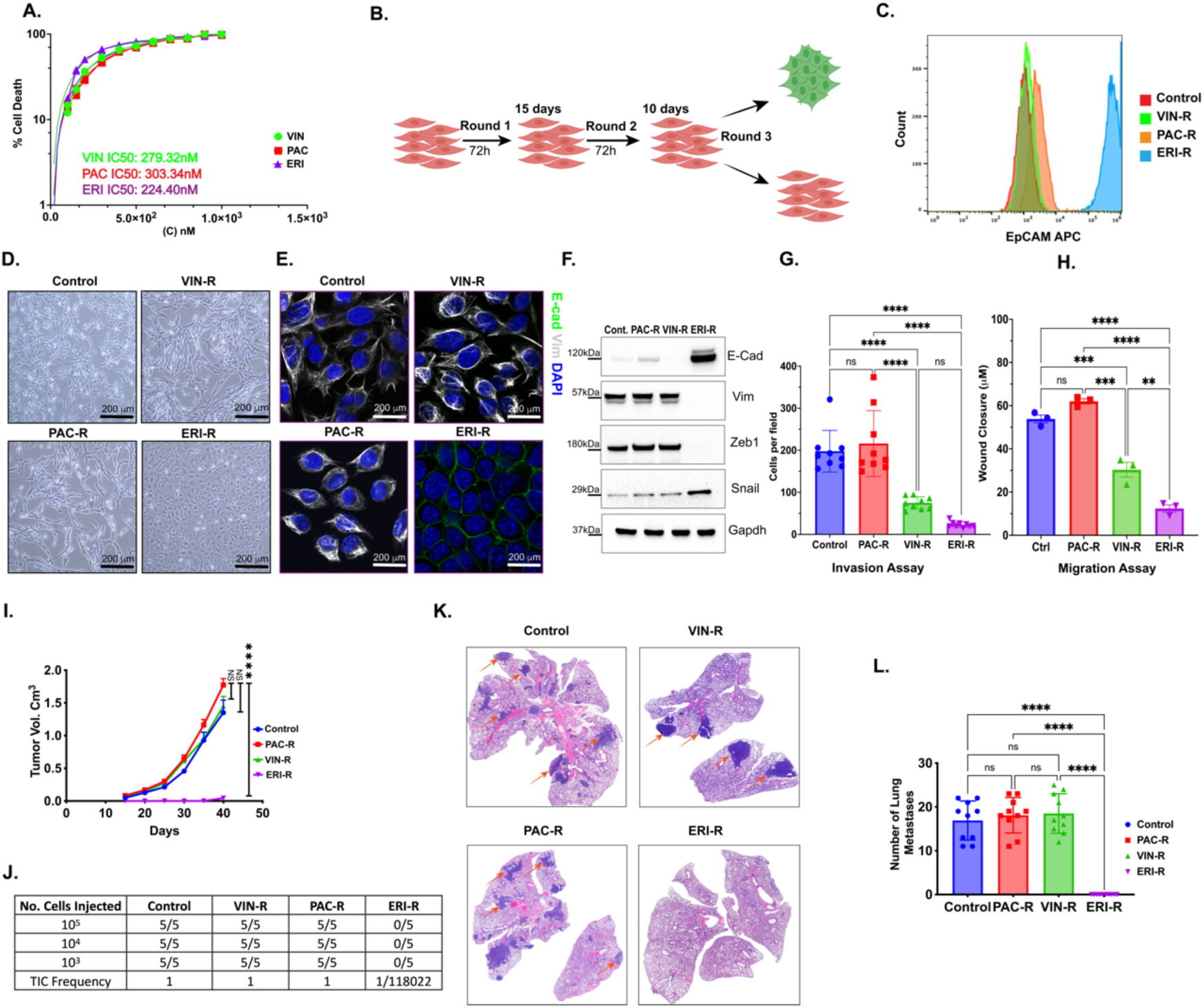
Eribulin induces MET in breast cancer cells. (A) Dose response curves in PB3 cells were generated to calculate IC_50_ values for eribulin, paclitaxel, and vinorelbine treatment. (B) Schematic representation of multi-round drug treatment procedure that resulted in the generation of resistant clones. (C) Histograms of FACS-based quantification of EpCAM expression in PB3 parental cells and resistant clones. Estimation of EMT state was carried out by (D) morphological assessment of brightfield images, (E) immunofluorescence of canonical markers E-cadherin (green) and Vimentin (white), and (F) immunoblotting to estimate protein levels of EMT markers in the PB3 parental line and resistant counterparts. Transwell invasion (G) and wound healing scratch assays (H) were carried out to assess the invasive properties of cells over 8 (invasion) and 16 (wound healing) hours. Data are shown as mean ± SD (n=3). The p value was determined by a two-way ANOVA. *p < 0.0001. Parental PB3 and resistant clones were transplanted orthotopically to assess tumor volume (I), tumor-initiating ability as assessed by limiting dilution transplantation (J) and metastatic ability of cells as observed by H&E staining (K) and enumeration (L). Data in (I), (L) are shown as mean ± SD (n=10). The p value was determined by a two-way ANOVA. *p < 0.0001.

Orthotopic implantation of PB3, VIN-R, and PAC-R cells into female NOD-scid IL2Rg^-/-^ (NSG) mice resulted in robust tumor growth. However, ERI-R cells did not readily form tumors (Fig. 1I) and exhibited a >100-fold reduction in tumor-initiating ability (Fig. 1J). Furthermore, ERI-R cells did not form spontaneous lung metastases despite the other cell lines yielding abundant lung lesions established from mammary fat pad implantation (Fig. 1K,L), likely a result of smaller primary tumor sizes. ERI-R MDA-MB-231 cells also exhibited loss of the cancer stem cell marker CD44 (Suppl. Fig. S1I), in line with the decreased tumor-initiating potential observed in ERI-R PB3 cells (Fig. 1I,J). These results confirm the ability of eribulin to induce MET in breast cancer cells, driving cells toward an epithelial state accompanied by a significant reduction in tumor-initiating potential.

### Clonal dynamics reveal a Lamarckian induction of epithelial-like chemotherapy-resistant cells

Drug treatment of a heterogeneous cell population results in a diverse array of fates that are determined by, amongst other factors, the starting transcriptional and epigenetic profiles of each cell. To elucidate the cellular trajectories that result from drug treatment, we developed an approach to <u>q</u>uery indexed <u>s</u>equences and <u>s</u>imultaneously <u>m</u>easure <u>e</u>pigenetic timelines (QISSMET). Similar approaches have been utilized to trace cell lineage through therapeutic bottlenecks ^9,10^. By analyzing changes in transcriptional and chromatin accessibility profiles induced by drug selection, QISSMET enables differentiation between pre-existing drug-resistant cells and emerging *de novo* cell types that acquire resistance. Changes that a given cell undergoes upon drug treatment would manifest as changes in gene expression and chromatin state, which would imply an epigenetic reprogramming leading to the emergence of a *de novo* cell state. Alternatively, if no changes in gene expression or chromatin state are observed, this would suggest that a selection event had occurred, amplifying a pre-existing drug-resistant population.

To characterize the nature of resistant cells following treatment with eribulin, we stably infected PB3 cells with an expressed barcode library. Barcoded cells were subject to 3-4 cycles of treatment with eribulin or paclitaxel (Fig. 2A), and an aliquot of cells was harvested before each cycle for single-cell RNA sequencing (scRNA-Seq). Eribulin treatment resulted in the emergence of cells (ERI3 and ERI4) that clustered distinctly from the parental population by transcriptomic analysis, while paclitaxel did not (Fig. 2B). Monocle pseudotime kinetics ^11–13^ revealed that cells undergoing eribulin treatment took multiple trajectories from the most primitive (parental) to the most advanced (ERI4) transcriptional state (Fig. 2C). In contrast, cells undergoing paclitaxel treatment took fewer distinct paths and remained more similar to the parental controls without uniform directionality (Fig. 2D).

**Figure 2:**
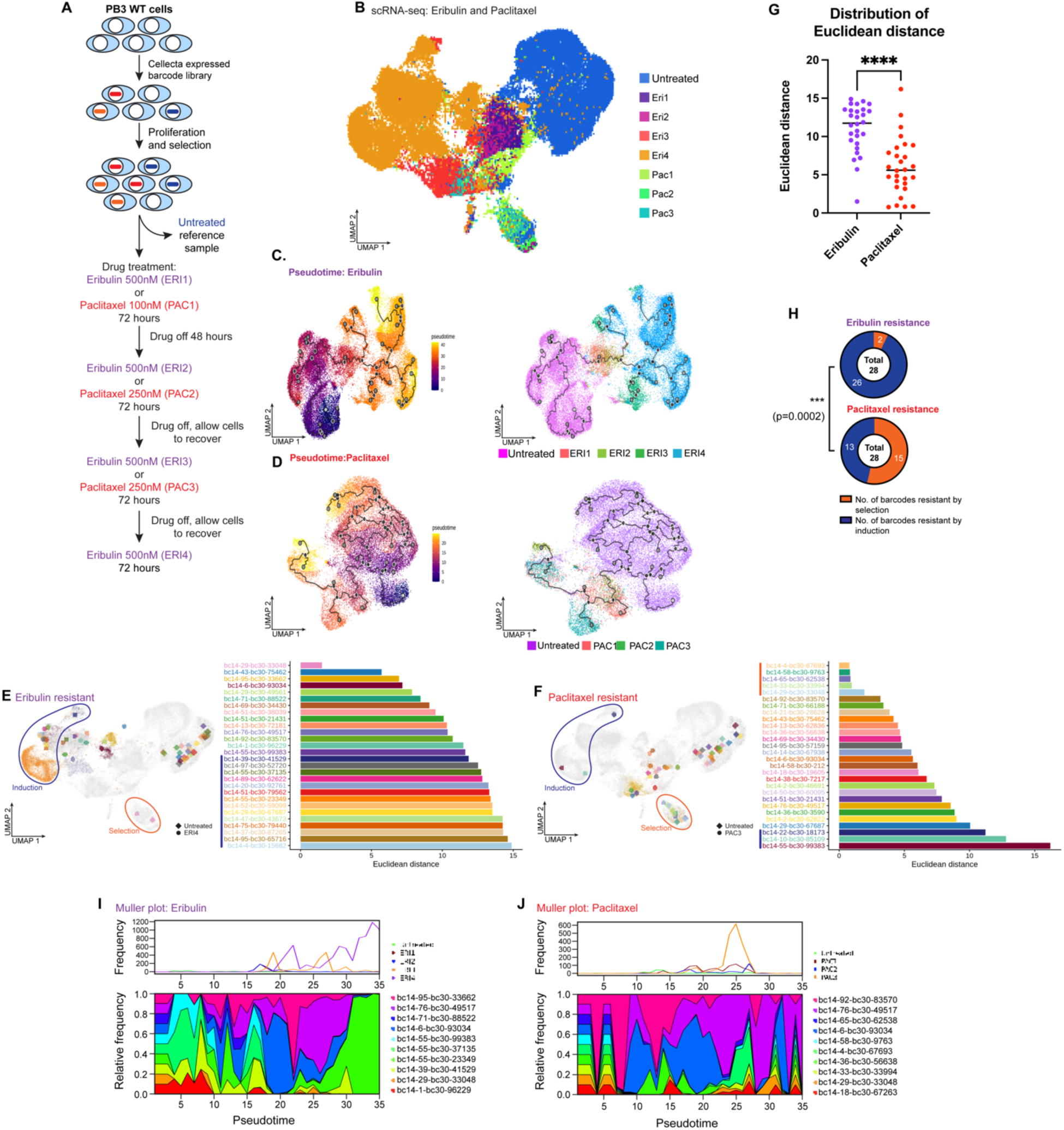
Clonal dynamics following drug treatment. (A) Schematic outlining lentiviral barcoding of cells and subsequent drug treatments strategy. (B) UMAP projection of scRNA-Seq data obtained from untreated PB3 and following treatment with either eribulin or paclitaxel. Monocle pseudotime projection showing the trajectory of tumor evolution upon treatment with (C) eribulin or (D) paclitaxel. The main lineage path or master path/node is indicated by the number 1 in the white circle. Sub-lineages from the master path (also called leaf nodes) are indicated by numbers in grey circles. Further sub-divisions or branches from the leaf nodes are indicated by numbers in black circles. Euclidean distance calculations between (E) ERI4 or (F) PAC3 and untreated cells with the same barcode. The UMAP overlay represent the median point of Eri4 or Pac3 and untreated cells. (G) Distribution of Euclidean distance values of eribulin-resistant and paclitaxel-resistant barcodes. Median for eribulin: 11.74, median for paclitaxel: 5.597. P<0.0001, Mann-Whitney test. (H) Proportions of induction vs selection resistance for eribulin and paclitaxel treatments. P=0.0002, Chi-Square test. Muller plot outlining the clonal diversity of the top 10 most abundant barcodes following (I) eribulin and (J) paclitaxel treatments, respectively.

We sought to determine whether eribulin treatment led to (i) an induction event that generated resistant cells with altered chromatin and transcriptional profiles, or (ii) the selection of pre-existing drug-resistant cells that were more similar to the parental population. We selected the 28 most highly enriched barcodes that contained ≥1 cell in the control and drug-treated clusters, and calculated their Euclidean distance based on median positions on UMAP plots. Drug-resistant cells with barcodes that had larger Euclidean distances resided in the far left (“Induction” region) of the UMAP (Fig. 2E), while drug-resistant cells with barcodes that had smaller Euclidean distance resided in the bottom island (“Selection” region) of the UMAP (Fig. 2F). Cells treated with eribulin vs. paclitaxel had significantly different Euclidian distances (median 11.74 vs. 5.59; Fig. 2G). Using a combination of Euclidian distance and Jaccard index, we calculated that ≥92% of barcodes in eribulin-resistant cells localized to the Induction region (Fig. 2H and Suppl. Fig. S2A-C), indicating that they underwent a shift in transcriptional profile as a result of eribulin treatment. In contrast, paclitaxel-resistant cells were transcriptionally more similar to the parental population with ≥50% of barcodes localizing to the Selection region (Fig. 2H and Suppl. Fig. S2B-E).

In addition to exhibiting altered transcriptional profiles, the resistant epithelial-like cell subpopulations that emerged from eribulin treatment also exhibited altered chromatin landscapes as observed by single-cell assay for transposase-accessible chromatin using sequencing (scATAC-Seq; Suppl. Fig. S2F,G). Plotting the evolutionary trajectories of cells as they underwent drug treatments revealed that both agents imposed severe bottlenecks that only a few barcodes (individual cells) overcame. While eribulin treatment resulted in the emergence of a dominant clonal subpopulation, clonal frequencies were preserved with less variation upon paclitaxel treatment (Fig. 2I,J). These results indicate that, in comparison to paclitaxel, eribulin treatment led to the evolution of resistant cells that were distinct from parental cells, consistent with their having undergone MET and reprogramming of transcriptional and chromatin states.

### Order of exposure to chemotherapeutics alters the efficacy

Current treatment strategies for TNBC often include the administration of neoadjuvant chemotherapy (NAC), which typically consists of a combination of an anthracycline (e.g., adriamycin) and cyclophosphamide followed by a taxane (e.g., paclitaxel) (AC-T) prior to surgical resection of the tumor. Patients with advanced/metastatic TNBC are treated with agents such as vinorelbine or eribulin, where eribulin is approved for use following prior treatment with ≥3 lines of chemotherapy that should have included an anthracycline and a taxane. We sought to understand differences in the ability of eribulin to induce MET when administered (a) in the treatment-naïve setting vs. (b) after exposure to other chemotherapeutics as occurs clinically. Data shown in Fig. 1C–F and Suppl. Fig. 1 indicated that primary treatment of TNBC cells with eribulin resulted in the generation of epithelial-like ERI-R cells. Upon subsequent secondary treatment of ERI-R cells with either paclitaxel or vinorelbine, >50% of cells became senescent and >40% of cells were eliminated via apoptosis (Fig. 3A,B and Suppl. Fig. S3A,B). In contrast, primary treatment with either paclitaxel or vinorelbine to generate PAC-R and VIN-R cells resulted in reduced sensitivity to secondary eribulin, with only ∼8% of cells undergoing senescence and ∼15% of cells undergoing apoptosis (Fig. 3C and Suppl. Fig. S3C,D).

**Figure 3:**
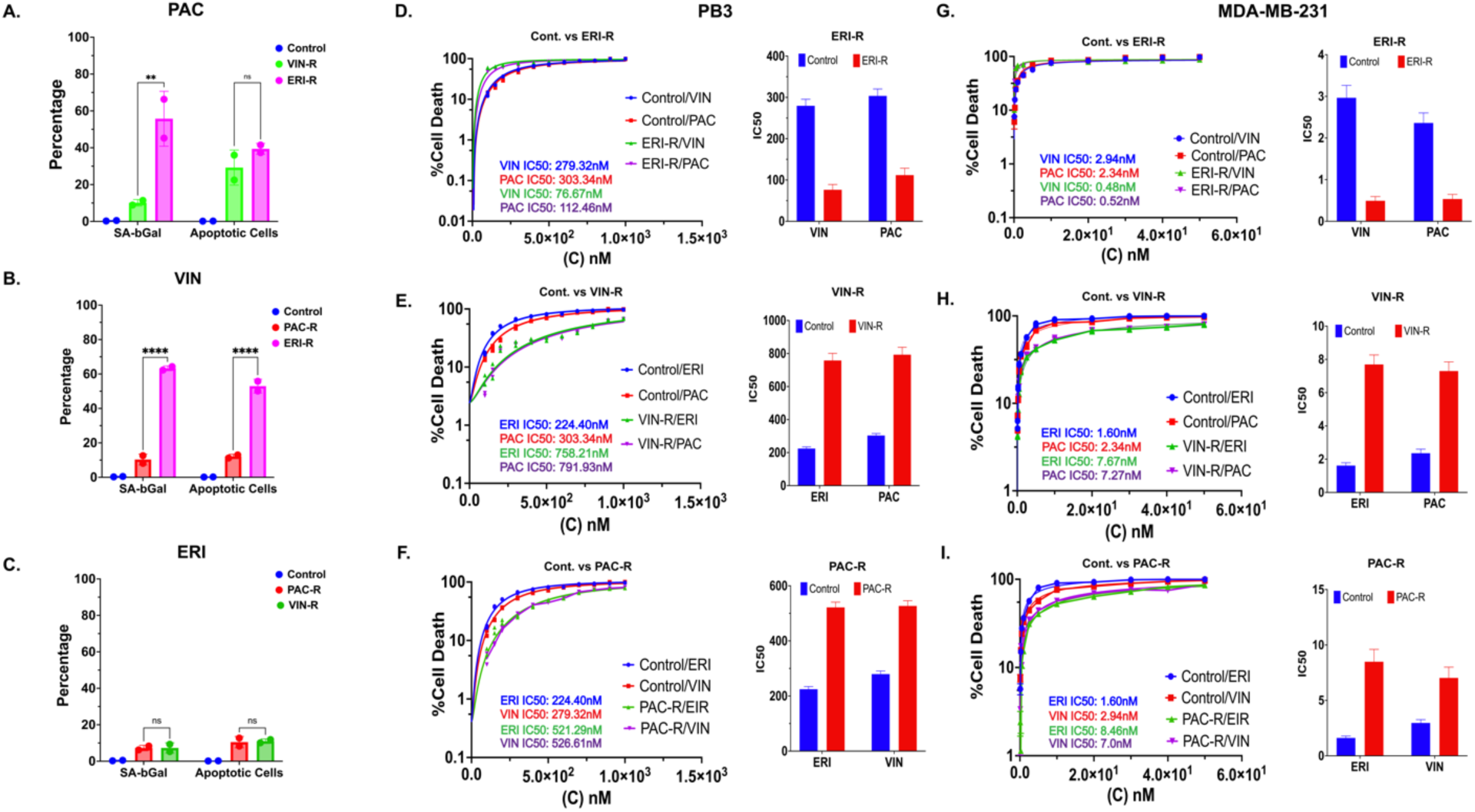
Eribulin pre-treatment induces sensitization to subsequent therapy. Senescence-associated beta-galactosidase assay and quantification of live-dead and apoptotic cells using SYTOX (flow cytometry) to evaluate drug response of PB3-derived (A) ERI-R and VIN-R cells to paclitaxel, (B) ERI-R and PAC-R cells to vinorelbine treatment, and (C) PAC-R and VIN-R cells to eribulin treatment. Dose-response curves to determine IC_50_ shift in PB3 parental cells and (D) ERI-R clone treated with paclitaxel and vinorelbine, (E) VIN-R clone treated with paclitaxel and eribulin, (F) PAC-R clone treated with various doses of paclitaxel and eribulin. Assays were also carried out in in MDA-MB-231 parental cells compared to (G) ERI-R clone treated with paclitaxel and vinorelbine, (H) VIN-R clone treated with paclitaxel and eribulin, (I) PAC-R clone treated with vinorelbine and eribulin (IC_50_ shift curves are shown on the left panel and IC_50_ shift quantification on the right panel).

We next sought to determine IC_50_ values of parental and drug-resistant PB3 cells to subsequent treatments with different chemotherapeutics. We treated PB3 and ERI-R cells with vinorelbine or paclitaxel, which resulted in a reduction in IC_50_ (i.e., increased sensitization) of ERI-R cells to vinorelbine (279.32 nM to 76 nM) compared to parental cells (a ∼3-fold reduction). Similarly, ERI-R cells were more sensitive to paclitaxel compared to parental cells (IC_50_ 112.46 nM vs. 303.34 nM; Fig. 3D). In contrast, VIN-R and PAC-R cells exhibited higher IC_50_ values to subsequent treatment with different chemotherapeutics (Fig. 3E,F). Similar patterns of chemosensitization and chemoresistance were observed in MDA-MB-231 cells (Fig. 3G–I). ERI-R MDA-MB-231 cells also exhibited widespread senescence upon treatment with vinorelbine or paclitaxel, whereas PAC-R and VIN-R cells did not (Suppl. Fig. S3E). These results point to MET induction with an agent such as eribulin being an efficient means of sensitizing therapy-naïve tumor cells to subsequent rounds of treatment with other agents.

### MET induction is accompanied by robust tumor regression and reduced metastatic burden

We next sought to understand how MET induction via eribulin treatment affected tumor growth and metastatic progression. We randomized autochthonous tumor-bearing MMTV-PyMT mice to treatment with vehicle, paclitaxel (20 mg/kg), vinorelbine (7 mg/kg) or eribulin (1.6 mg/kg) twice weekly for 2 weeks. While vinorelbine and paclitaxel each induced partial tumor regression or stasis, eribulin led to robust tumor regression in all cases (Fig. 4A). Furthermore, eribulin treatment induced a near-complete inhibition of metastatic growth in the lungs that was significantly more effective than other drugs (Fig. 4B). In contrast to the vinorelbine or paclitaxel treatments that resulted in high-grade poorly differentiated tumors containing an abundance of vimentin-expressing (i.e., mesenchymal-like) tumor cells, eribulin treatment resulted in low-to moderate-grade well-differentiated tumors containing islands of malignant cells primarily expressing the epithelial marker E-cadherin (Fig. 4C and Suppl. Fig. S4A).

**Figure 4:**
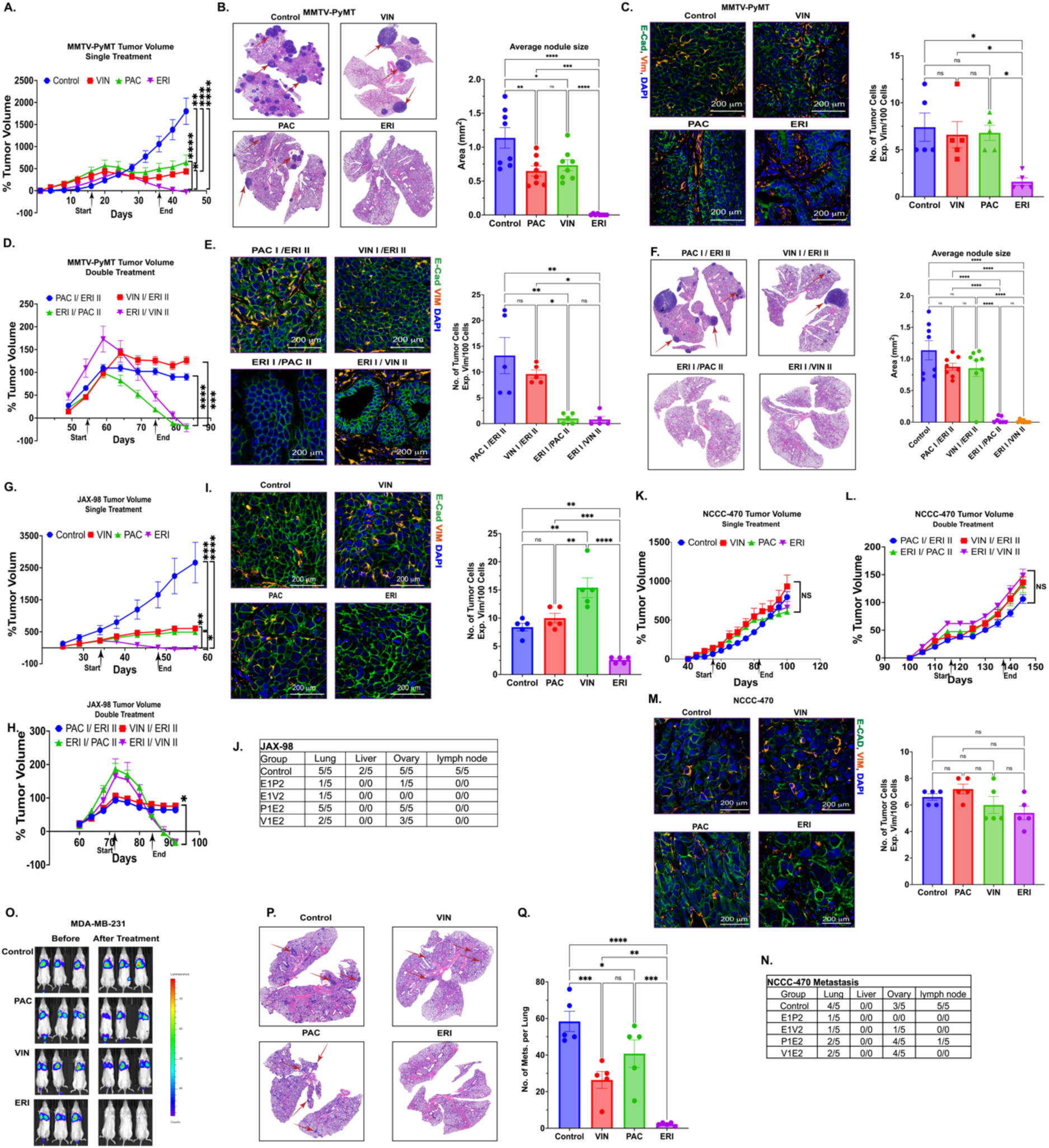
MET induction is accompanied by robust tumor regression and reduced metastatic burden. MMTV-PyMT tumor-bearing mice were treated with vinorelbine, paclitaxel, and eribulin to assess (A) tumor growth rate, (B) tumor metastasis to the lung (Arrows indicate metastatic lesions) and (C) EMT status using immunohistofluorescence of canonical EMT markers E-cadherin (green) and Vimentin (red), and The residual MMTV-PyMT tumor cells were treated with a second round of therapy to assess (D) tumor growth rate, (E) EMT status using immunofluorescence of canonical markers E-cadherin (green) and Vimentin (red), in addition to (F) tumor metastasis to the lung (Arrows indicate metastatic lesions). Two triple negative breast cancer-derived PDXs, the naïve JAX-98 and pretreated NCCC-470 were treated with vinorelbine, paclitaxel, and eribulin to evaluate the effect of treatment on, (G and K) tumor growth rate in single drug treatment regimens, (I and M) EMT status using immunohistofluorescence of canonical markers E-cadherin (green) and Vimentin (red), (H and L) tumor growth rate, and (J and N) distribution patterns of tumor metastasis from the sequential double treatment regimens. To study the effect of vinorelbine, paclitaxel, and eribulin on already-established metastasis, (O) MDA-MB-231 cells were injected through tail vein and tumor growth examined using bioluminescent imaging before and after treatments in addition to (P and Q) H&E staining and quantification of metastatic lesions following treatment. (∗p <0.05, ∗∗p < 0.001, ∗∗∗p > 0.0005, ∗∗∗∗p < 0.0001, Two-Way ANOVA, n=5).

To determine the effects of prior chemotherapeutic exposure on sensitivity to further chemotherapeutics, MMTV-PyMT tumor-bearing mice received primary treatment with 5 doses of paclitaxel or vinorelbine over 2 weeks, followed by a 2-week drug holiday and secondary treatment with eribulin for 2 weeks. We also carried out similar treatments with primary eribulin followed by secondary paclitaxel or vinorelbine. Tumors in mice that received primary eribulin underwent robust regression in response to secondary paclitaxel or vinorelbine (Fig. 4D). In contrast, secondary eribulin only induced stasis in tumors that received primary vinorelbine or paclitaxel. Moreover, tumors harvested following primary eribulin and secondary paclitaxel or vinorelbine contained higher proportions of E-cadherin-expressing malignant cells compared to tumors that received eribulin as secondary treatment (Fig. 4E). These observed differences in tumor growth and composition also resulted in differences in lung metastases: primary eribulin induced near-complete abrogation of metastatic propensity (Fig. 4F).

We then tested the effects of chemotherapeutics in 4 TNBC patient-derived xenograft (PDX) models: 2 from treatment-naïve patients (NCI-140, JAX-98) and 2 from patients treated with neoadjuvant chemotherapy (NCCC-470, JAX-91). In the treatment-naïve JAX-98 tumors, drug treatment effects were similar to those observed in MMTV-PyMT tumors. Eribulin induced near-complete regression of JAX-98 tumors (Fig. 4G–H), yielding residual lesions that were well-differentiated with an abundance of E-cadherin-expressing cells (Fig. 4I and Suppl. Fig. S4B). Mice that received primary eribulin had no metastases across various organs (lung, liver, ovary, lymph node). In contrast, vehicle-treated mice and those that received primary vinorelbine or paclitaxel followed by secondary eribulin exhibited significant metastatic burden across organs (Fig. 4J). Similar observations were made in a second treatment-naïve PDX model where eribulin induced near-complete regression of tumors, yielding residual lesions that were well-differentiated with an abundance of E-cadherin-expressing cells and enlarged senescent-like cells (NCI-140; Suppl. Fig. S4D-H). In contrast, growth of NCCC-470 tumors was not significantly affected by any chemotherapeutic regimen tested (Fig. 4K/L), with all tumors retaining poor differentiation status (Suppl. Fig. S4C), containing abundant vimentin-expressing cells, and generating metastases in lungs and ovaries (Fig. 4M/N). Similar observations were made in the JAX-91 model derived from a patient after neoadjuvant chemotherapy (Suppl. Fig. S4I-M).

Metastasis and EMT are early events in the tumorigenic process, and patients presenting with early-stage breast cancer often already have disseminated cancer cells in other organs ^14,15^. It is thus imperative in preclinical therapeutic studies to assess effects on metastases. We injected luciferase-expressing MDA-MB-231 cells via tail vein to establish lung metastases. Once metastatic colonies were detected by bioluminescence imaging, mice were treated with 4 rounds of vehicle, paclitaxel, vinorelbine, or eribulin. While vinorelbine and paclitaxel slowed (or did not affect) the growth of lung metastases, eribulin induced near-complete regression (Fig. 4O–Q). These data provide evidence of the ability of eribulin treatment to induce the regression of already established metastases, presumably via induction of MET or a combination of its MET-inducing and anti-proliferative effects.

### MET induction reduces epithelial-mesenchymal heterogeneity in tumors

Eribulin pre-treatment increased proportions of E-cadherin-expressing cells and decreased vimentin-expressing cells in TNBC PDX models (Fig. 4I,M and Suppl. Fig. S4E,J). To quantify EMT state and the extent of epithelial-mesenchymal heterogeneity (EMH) in tumors before and after drug treatment, we employed a multi-round, multiplexed immunostaining protocol. This entropy-based analytic method quantifies EMH and the extent of mesenchymal traits (EMT), metrics that predict patient survival ^16^. Treatment of mice bearing treatment-naïve NCI-140 tumors with eribulin reduced proportions of cells with an intermediate EMT state (triple-positive for E-cadherin, Keratin 8, and vimentin) accompanied by an increased representation of keratin 14-expressing epithelial-like cells (Fig. 5A,G). In contrast, eribulin was unable to shift the representation of cell subpopulations in NCCC-470 tumors derived from a patient treated with neoadjuvant chemotherapy (Fig. 5B,H).

**Figure 5:**
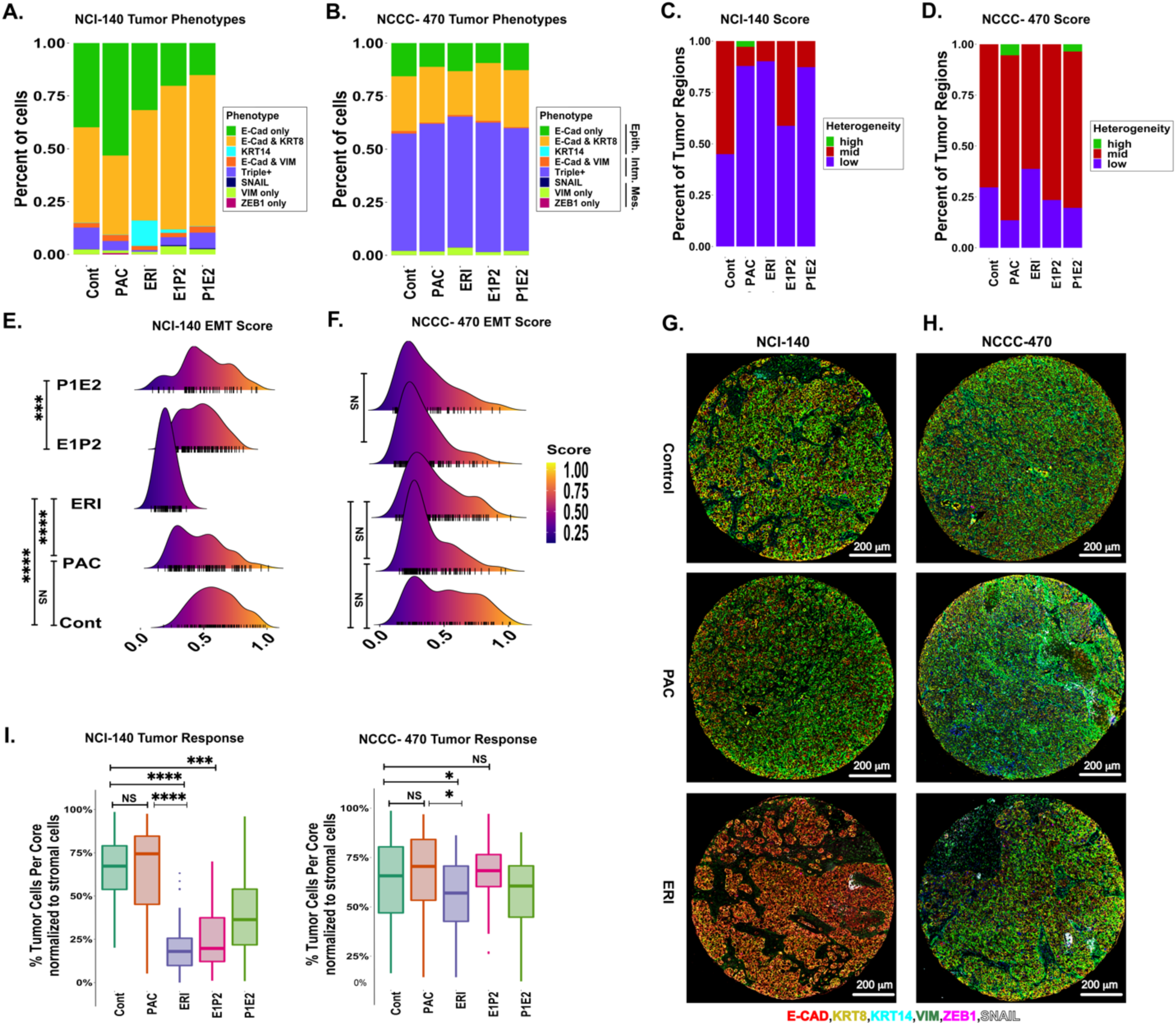
MET induction leads to the reduction in epithelial-mesenchymal heterogeneity of tumors. Multiplexed, multi-round TSA staining of EMT markers in triple negative breast cancer derived PDXs (NCI-140 and NCCC-470) treated with paclitaxel and eribulin as a single and sequential double treatment to evaluate (A and B) distribution of tumor phenotypes by calculating the percent of tumor cells expressing different EMT markers, (C and D) entropy-based tumor heterogeneity levels, (E and F) EMT score (Five tumors and >100 ROI/per tumor were analyzed). (G) Representative examples of multi-colored TSA staining of naïve NCI-140 and (I) pretreated NCCC-470 PDX tumors treated with paclitaxel and eribulin. Tumors were stained using anti-E-CAD (red), anti-KRT8 (yellow), anti-KRT14 (cyan), anti-VIM (green), anti-ZEB1 (magenta), and SNAIL (white) antibodies and counterstained with DAPI for the nuclei (blue). (G) Calculation of Tumor Response to Drug Treatment. (∗p <0.05, ∗∗p < 0.001, ∗∗∗p > 0.0005, ∗∗∗∗p < 0.0001, Two-Way ANOVA, (n=5)).

Tumors were classified into low, moderate (mid), and high EMH based on the extent of cellular diversity along the E-M axis. Eribulin or paclitaxel decreased the frequency of tumor regions containing moderate levels of EMH while increasing regions containing low EMH (Fig. 5C). Paclitaxel also induced focal increases to yield EMH-high regions. Similar observations were made in NCCC-470 tumors, but the extent of changes in EMH were more modest (Fig. 5D).

EMT scoring was also used to score tumor status along the E-M axis with 0 being most epithelial and 1 being most mesenchymal. Eribulin induced an epithelial-oriented shift in mean EMT score from 0.57 in control-treated NCI-140 (treatment-naïve) tumors to 0.16 (Fig. 5E). In contrast, NCCC-470 tumors did not exhibit a shift in EMT score upon drug treatment (Fig. 5F), in line with resistance to eribulin-induced MET (Fig. 4M).

We also quantified tumor response to therapy based on changes in proportions of cancer vs. stromal cells ^16^. Single-agent eribulin and paclitaxel each reduced proportions of cancer cells in NCI-140 tumors by means of 80% and 35%, respectively (Fig. 5I). In contrast, NCCC-470 tumors exhibited weaker responses to treatments. Similar comparisons were made with another set of treatment-naïve (JAX-98) and pre-treated (JAX-91) PDX models, in which eribulin treatment resulted in varied phenotypes (Suppl. Fig. S5A), heterogeneity (Suppl. Fig. S5B) and EMT score levels (Suppl. Fig. S5C) depending on pretreatment status. Importantly, the tumor response criteria were reproducibly higher when treatment-naïve tumors, but not pretreated tumors received eribulin (Suppl. Fig. S5D/E). These data reinforce our earlier observations that the effects of eribulin on MET induction and tumor regression are most profound when administered in the treatment-naïve setting.

### Treatment with eribulin induces a shift in the chromatin profiles of cancer cells

In light of the observed functional properties of eribulin, we sought to further explore its mechanism of action that contributes to its ability to induce an MET and associated sensitization to treatment by cytotoxic agents. Treatment of PB3 cells with eribulin led to the emergence of ERI-R cells that exhibited a significantly altered transcriptional profile relative to parental cells and PAC-R and VIN-R counterparts (Fig. 6A,B; Suppl. Fig. S6A). Gene set enrichment analysis revealed significant enrichment for hallmark gene sets involved in EMT, the majority of which were downregulated in ERI-R cells compared to parental controls (Fig. 6C/D).

**Figure 6:**
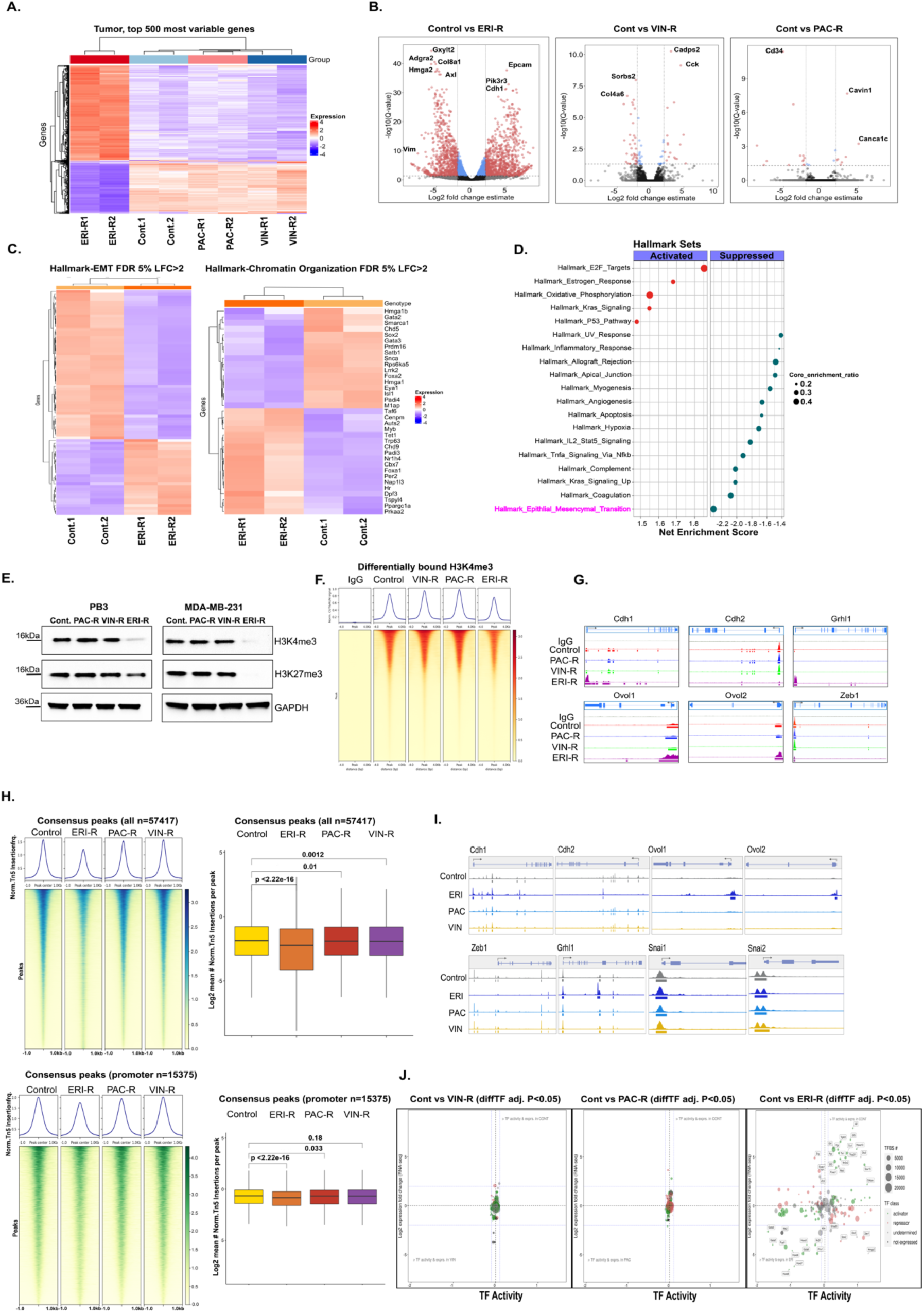
Treatment with eribulin induces a shift in the chromatin profile of cancer cells. Gene expression analysis differences in PB3 parental, VIN-R, PAC-R, and ERI-R cells were illustrated by (A) heat map of the top 500 differentially expressed genes and (B) volcano plot of all genes significantly regulated upon treatment (log2-fold-change indicates the mean expression level for each gene, Benjamini-Hochberg corrected p-value threshold = 0.01). (C) Heatmap of EMT and chromatin organization gene set enrichment in PB3 parental and ERI-R cells as well as (D) dot plot outlining hallmark sets enrichment. The size of the circle depicts the significance of the pathways, red dots indicate activated pathways, green dots indicate suppressed pathways. (E) Immunoblotting showing levels of H3K4me3 and H3K27me3 evaluated in PB3 and MDA-MB-231 parental and resistant clones and (F) the average CUT&RUN enrichment profile of H3K4me3 in PB3 control and resistant clones treated with the indicated drugs (top). Heatmap illustrating the CUT&RUN signal 4 kb up-and downstream of the different peaks. (G) H3K4me3 localization at genomic loci of canonical EMT genes was evaluated by CUT&RUN signal track analysis. ATAC-sequencing was performed in PB3 parental and resistant clones to explore chromatin accessibility at different regions (H) peak accessibility surrounding all consensus regions (top) and promoter-associated regions (down) and (I) peak accessibility of different EMT markers across all conditions. TF motifs are highlighted below each signal track. (J) ATAC-seq and RNA-seq integration analysis. Advanced volcano plot of highly significant transcription factors determined by diffTF from ATAC-seq along the X-axis and Log2 fold gene expression values of transcription factors on the y-axis. Transcription factor classification is displayed by bubble color, and number of transcription factor binding sites used to determine TF activity is plotted as bubble size.

Given the observed significant changes in transcriptional programs with eribulin selection, we reasoned that these changes resulted from alterations to the chromatin state of cancer cells. Indeed, the levels of the H3K4me3 activation mark and the H3K27me3 repressive mark were reduced in ERI-R derivatives, but not PAC-R or VIN-R derivatives, from PB3, MDA-MB-231 and SUM159 cells (Fig. 6E and Suppl. Fig. S6B). H3K4me3 cistrome profiling revealed overall reduced representation of this activating histone mark in ERI-R PB3 cells (Fig. 6F). However, this reduction in H3K4me3-DNA binding was not universal, with some genomic loci coding for epithelial markers such as *Cdh1*, *Grhl1*, *Ovol1* and *Ovol2* gaining this mark in promoter regions in ERI-R cells, but not PAC-R or VIN-R cells (Fig. 6G).

Assessing the chromatin accessibility landscape of cells by ATAC-Seq also revealed an overall more closed chromatin profile with fewer and different accessible regions of chromatin in ERI-R derivatives compared to parental PB3 cells (Fig. 6H; Suppl. Fig. S6C-D). Consistent with H3K4me3 profiling, chromatin surrounding loci encoding epithelial markers (*Cdh1*, *Ovol1*, *Ovol2*) was more accessible in ERI-R cells (Fig. 6I). Integrated analysis of RNA-seq and ATAC-seq data with DiffTF ^17^ inferred large-scale changes in the activity of key transcription factors in ERI-R cells compared to PB3 controls, but not in VIN-R or PAC-R cells (Fig. 6J and Suppl. Fig. S6E/F). These data collectively point to eribulin treatment resulting in more pronounced reprogramming of the chromatin and transcriptional state of TNBC cells compared to treatment with other microtubule dynamics inhibitors, suggesting alternate and distinct mechanisms of drug action.

### ZEB1-SWI/SNF interactions are required for maintenance of mesenchymal state

To identify protein targets involved in chromatin/transcriptional responses to eribulin, we conducted an unbiased proteome integral solubility alteration (PISA) assay ^18^ to identify proteins that exhibit altered thermal stability within 4 h in the presence of eribulin in PB3 cells. Among 8000 proteins detected in all samples, 40 proteins had significant (p<0.05) alterations in thermal stability from eribulin treatment (Table 1 and Suppl. Table S1). We focused validation studies on proteins with DSm≥0.5 and –Log^10^≥1.5 with known roles in chromatin regulation. Smrd1 (*Smarcd1*) and Smrd3 (*Smarcd3*) are SWI/SNF family ATP-dependent chromatin remodelers that exhibited thermal stability shifts with eribulin treatment, indicating that they may be bound by the drug (Fig. 7A and Suppl. Fig. S7A). CRISPR/Cas9-mediated knockout of *Smarcd1*, *Smarcd2*, or *Smarcd3* in PB3 cells reduced spindle-shaped morphology, reduced Zeb1, and increased E-cadherin (Fig. 7B and Suppl. Fig. S7B/C). Triple knockout of *Smarcd1*/*2*/*3* phenocopied eribulin treatment, shifting PB3 and MDA-MB-231 cells towards an epithelial state (Fig. 7B and Suppl. Fig. S7F/G), and mirrored eribulin-induced decreases in H3K4me3 and H3K27me3 (Fig. 7C). Rescue through human *SMARCD1* expression in *Smarcd1*-knockout PB3 cells suppressed E-cadherin levels and restored Zeb1, indicating that SMARCD1 signaling drives a mesenchymal state (Suppl. Fig. S7D).

**Figure 7:**
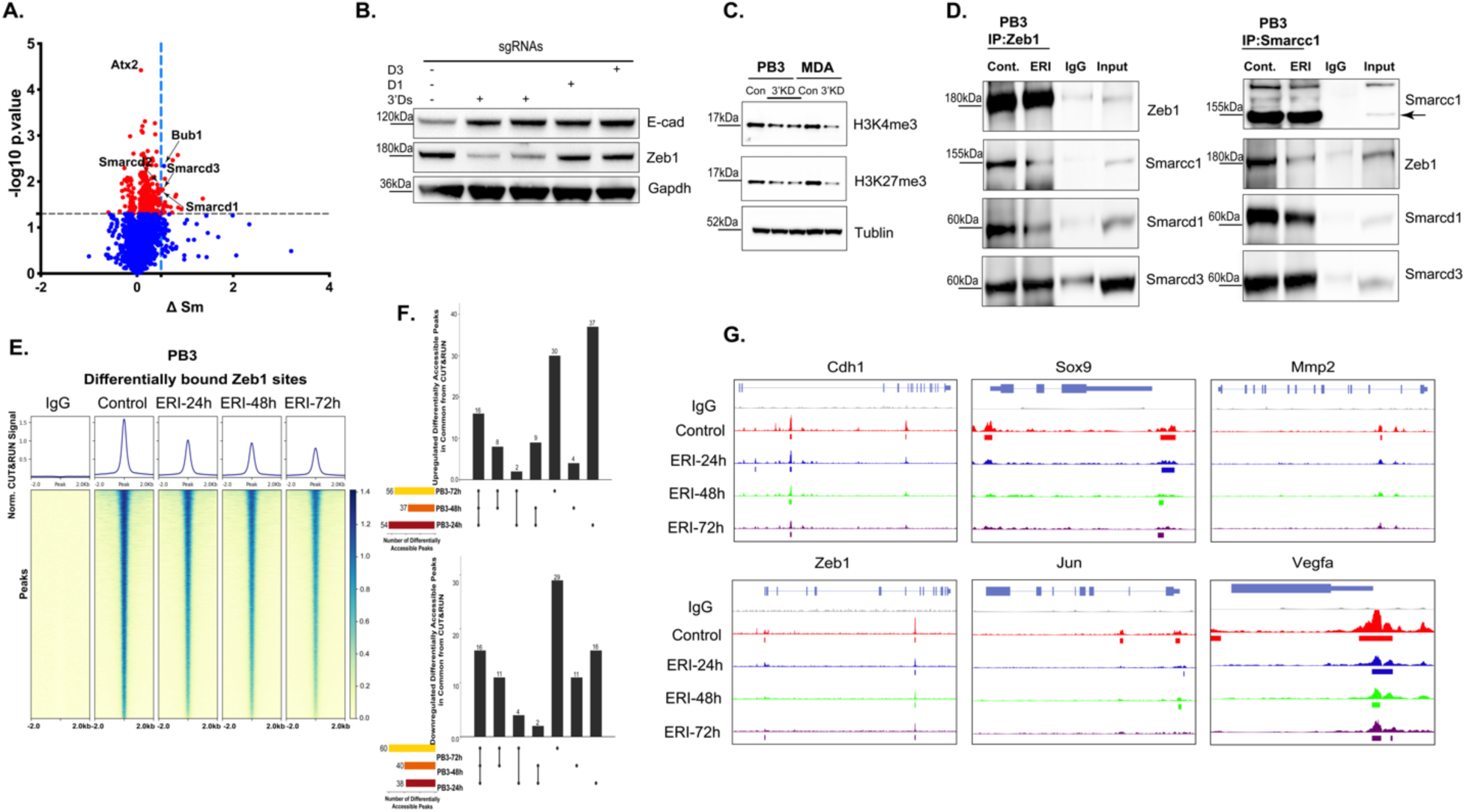
ZEB1-SWI/SNF interactions are necessary for maintenance of a mesenchymal state. (A) Proteome Integral Solubility Alteration of PB3 cells treated with 300nM eribulin for 4 hours to explore the non-tubulin targets of eribulin. Indicated proteins show the highest delta Sm and p-value. (Delta Sm > 0.5 and p < 0.05, T-test). CRISPR/Cas9-mediated deletion of Smarcd1, Smarcd2, and Smarcd3 genes and estimation of EMT state as carried out by (B) immunoblotting to estimate protein levels of EMT markers in the PB3 parental line and resistant counterparts. (C) Immunoblotting to evaluate levels of histone marks H3K4me3 and H3K27me3 in PB3 and MDA-MB-231 parental cells and upon combined knockout of Smarcd1, 2 and 3. The effect of eribulin on the Zeb1-SWI/SNF interaction was assessed by (D) co-immunoprecipitation using an anti-Zeb1 antibody (left panel) and anti-Smarcc1 antibody and probed for indicated proteins (right panel) in PB3 parental and eribulin-treated cells. The genome-wide occupancy of Zeb1 binding sites in PB3 parental cells and those treated with eribulin for 24, 48, and 72 hours was evaluated using CUT&RUN (E). The average CUT&RUN enrichment profile (top) and heatmap illustrating the CUT&RUN signal 2 kb up-and downstream of the different peaks. (F) UpSet plots of differentially accessible peaks at the indicated time points and (G) signal tracks of Zeb1 localization at target gene loci during eribulin treatment.

**Table 1.**
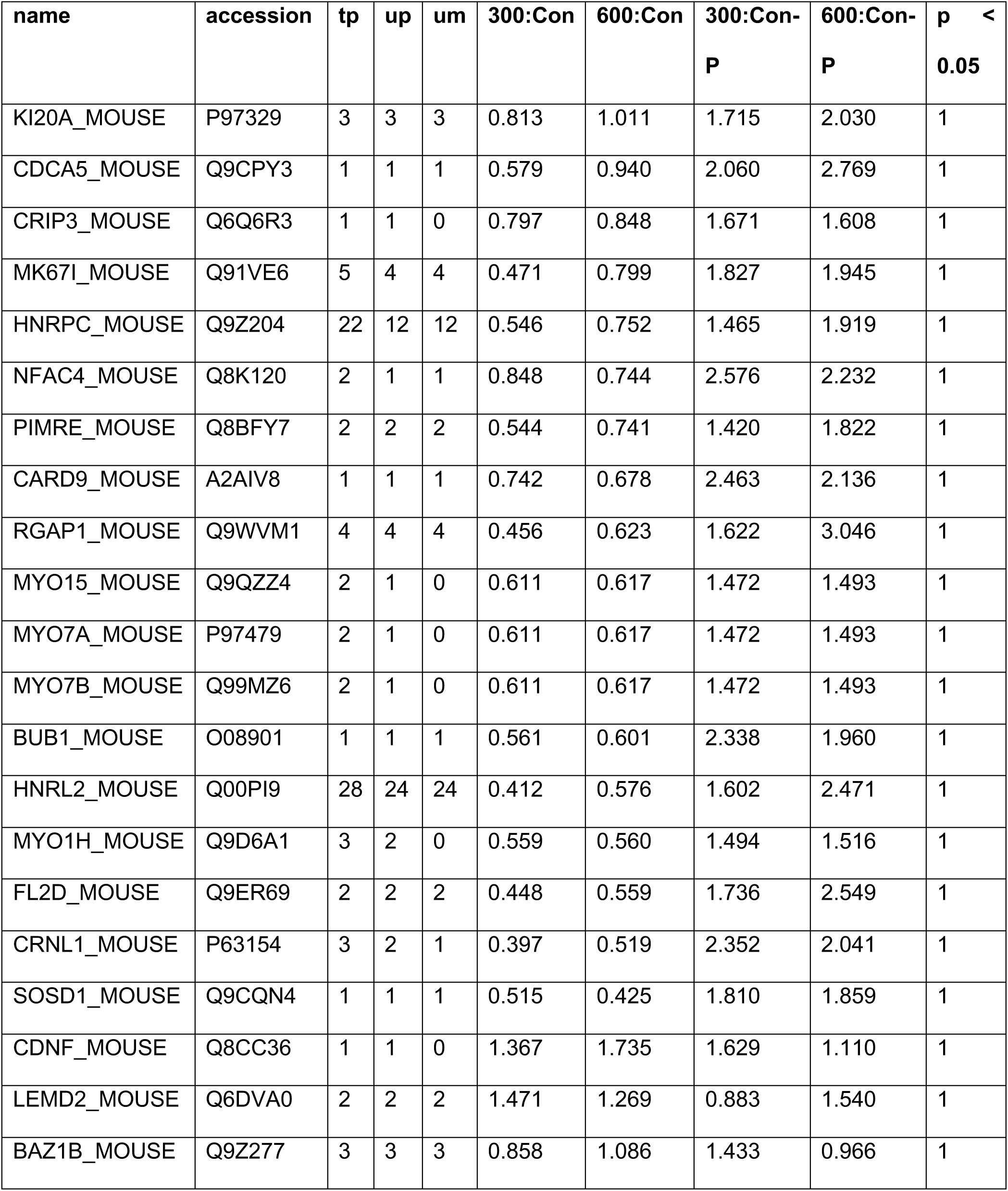
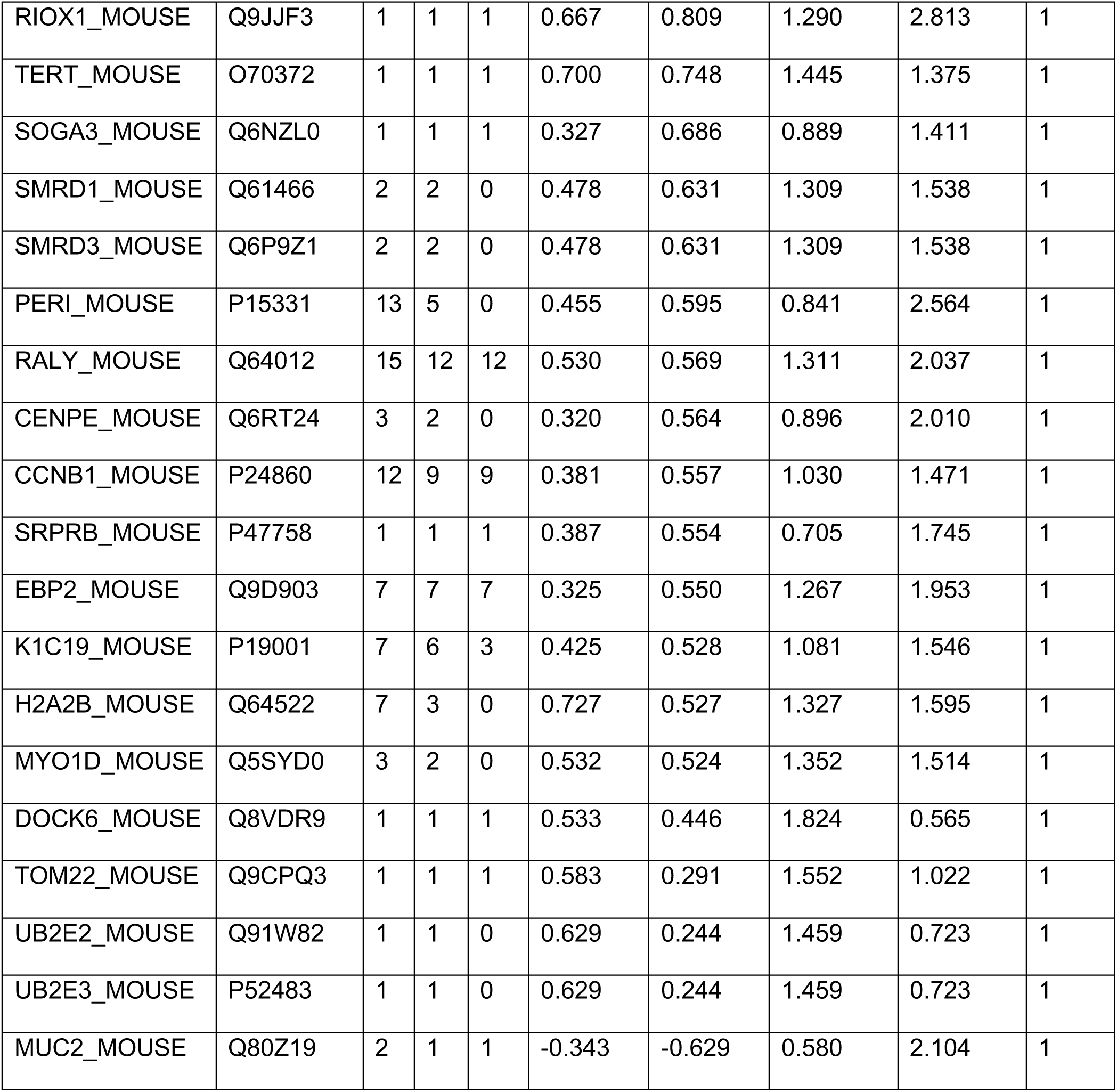
PISA Significant Results

| name | accession | tp | up | um | 300:Con | 600:Con | 300:Con-P | 600:Con-P | p <<br>0.05 |
| --- | --- | --- | --- | --- | --- | --- | --- | --- | --- |
| KI20A_MOUSE | P97329 | 3 | 3 | 3 | 0.813 | 1.011 | 1.715 | 2.030 | 1 |
| CDCA5_MOUSE | Q9CPY3 | 1 | 1 | 1 | 0.579 | 0.940 | 2.060 | 2.769 | 1 |
| CRIP3_MOUSE | Q6Q6R3 | 1 | 1 | 0 | 0.797 | 0.848 | 1.671 | 1.608 | 1 |
| MK67I_MOUSE | Q91VE6 | 5 | 4 | 4 | 0.471 | 0.799 | 1.827 | 1.945 | 1 |
| HNRPC_MOUSE | Q9Z204 | 22 | 12 | 12 | 0.546 | 0.752 | 1.465 | 1.919 | 1 |
| NFAC4_MOUSE | Q8K120 | 2 | 1 | 1 | 0.848 | 0.744 | 2.576 | 2.232 | 1 |
| PIMRE_MOUSE | Q8BFY7 | 2 | 2 | 2 | 0.544 | 0.741 | 1.420 | 1.822 | 1 |
| CARD9_MOUSE | A2AIV8 | 1 | 1 | 1 | 0.742 | 0.678 | 2.463 | 2.136 | 1 |
| RGAP1_MOUSE | Q9WVM1 | 4 | 4 | 4 | 0.456 | 0.623 | 1.622 | 3.046 | 1 |
| MYO15_MOUSE | Q9QZZ4 | 2 | 1 | 0 | 0.611 | 0.617 | 1.472 | 1.493 | 1 |
| MYO7A_MOUSE | P97479 | 2 | 1 | 0 | 0.611 | 0.617 | 1.472 | 1.493 | 1 |
| MYO7B_MOUSE | Q99MZ6 | 2 | 1 | 0 | 0.611 | 0.617 | 1.472 | 1.493 | 1 |
| BUB1_MOUSE | O08901 | 1 | 1 | 1 | 0.561 | 0.601 | 2.338 | 1.960 | 1 |
| HNRL2_MOUSE | Q00PI9 | 28 | 24 | 24 | 0.412 | 0.576 | 1.602 | 2.471 | 1 |
| MYO1H_MOUSE | Q9D6A1 | 3 | 2 | 0 | 0.559 | 0.560 | 1.494 | 1.516 | 1 |
| FL2D_MOUSE | Q9ER69 | 2 | 2 | 2 | 0.448 | 0.559 | 1.736 | 2.549 | 1 |
| CRNL1_MOUSE | P63154 | 3 | 2 | 1 | 0.397 | 0.519 | 2.352 | 2.041 | 1 |
| SOSD1_MOUSE | Q9CQN4 | 1 | 1 | 1 | 0.515 | 0.425 | 1.810 | 1.859 | 1 |
| CDNF_MOUSE | Q8CC36 | 1 | 1 | 0 | 1.367 | 1.735 | 1.629 | 1.110 | 1 |
| LEMD2_MOUSE | Q6DVA0 | 2 | 2 | 2 | 1.471 | 1.269 | 0.883 | 1.540 | 1 |
| BAZ1B_MOUSE | Q9Z277 | 3 | 3 | 3 | 0.858 | 1.086 | 1.433 | 0.966 | 1 |
| RIOX1_MOUSE | Q9JJF3 | 1 | 1 | 1 | 0.667 | 0.809 | 1.290 | 2.813 | 1 |
| TERT_MOUSE | O70372 | 1 | 1 | 1 | 0.700 | 0.748 | 1.445 | 1.375 | 1 |
| SOGA3_MOUSE | Q6NZL0 | 1 | 1 | 1 | 0.327 | 0.686 | 0.889 | 1.411 | 1 |
| SMRD1_MOUSE | Q61466 | 2 | 2 | 0 | 0.478 | 0.631 | 1.309 | 1.538 | 1 |
| SMRD3_MOUSE | Q6P9Z1 | 2 | 2 | 0 | 0.478 | 0.631 | 1.309 | 1.538 | 1 |
| PERI_MOUSE | P15331 | 13 | 5 | 0 | 0.455 | 0.595 | 0.841 | 2.564 | 1 |
| RALY_MOUSE | Q64012 | 15 | 12 | 12 | 0.530 | 0.569 | 1.311 | 2.037 | 1 |
| CENPE_MOUSE | Q6RT24 | 3 | 2 | 0 | 0.320 | 0.564 | 0.896 | 2.010 | 1 |
| CCNB1_MOUSE | P24860 | 12 | 9 | 9 | 0.381 | 0.557 | 1.030 | 1.471 | 1 |
| SRPRB_MOUSE | P47758 | 1 | 1 | 1 | 0.387 | 0.554 | 0.705 | 1.745 | 1 |
| EBP2_MOUSE | Q9D903 | 7 | 7 | 7 | 0.325 | 0.550 | 1.267 | 1.953 | 1 |
| K1C19_MOUSE | P19001 | 7 | 6 | 3 | 0.425 | 0.528 | 1.081 | 1.546 | 1 |
| H2A2B_MOUSE | Q64522 | 7 | 3 | 0 | 0.727 | 0.527 | 1.327 | 1.595 | 1 |
| MYO1D_MOUSE | Q5SYD0 | 3 | 2 | 0 | 0.532 | 0.524 | 1.352 | 1.514 | 1 |
| DOCK6_MOUSE | Q8VDR9 | 1 | 1 | 1 | 0.533 | 0.446 | 1.824 | 0.565 | 1 |
| TOM22_MOUSE | Q9CPQ3 | 1 | 1 | 1 | 0.583 | 0.291 | 1.552 | 1.022 | 1 |
| UB2E2_MOUSE | Q91W82 | 1 | 1 | 0 | 0.629 | 0.244 | 1.459 | 0.723 | 1 |
| UB2E3_MOUSE | P52483 | 1 | 1 | 0 | 0.629 | 0.244 | 1.459 | 0.723 | 1 |
| MUC2_MOUSE | Q80Z19 | 2 | 1 | 1 | -0.343 | -0.629 | 0.580 | 2.104 | 1 |

Previous work has reported that members of the SWI/SNF family can interact with ZEB1, enabling repression of E-cadherin (*CDH1*) transcription and inducing an EMT ^19^. We hypothesized that the specificity of SMARCD proteins for EMT traits could be a result of interaction with ZEB1 to modulate ZEB1 function. Indeed, we observed Zeb1/ZEB1 interactions with Smarcd1/SMARCD1, Smarcd3/SMARCD3 and Smarcc1/SMARCC1 in PB3 and MDA-MB-231 cells (Fig. 7D and Suppl. Fig. S7H). These interactions were inhibited by treatment with eribulin.

We tested the effects of interactions between Zeb1/Smarcd1 and Zeb1/Smarcc1 on DNA binding and transcriptional activity. In PB3 and MDA-MB-231 cells, eribulin progressively reduced Zeb1/ZEB1 binding to DNA across the genome (Fig. 7E,F and Suppl. Fig. S7I,J). Eribulin elicited a time-dependent reduction in Zeb1 binding to the promoter regions of target genes *Cdh1* and *Mmp2* (Fig. 7G and Suppl. Fig. S7K). These results point to a role for Smarcd1 and Smarcc1 as essential co-factors maintain the ability of Zeb1 to maintain a mesenchymal state.

### Eribulin reduces EMT in primary human triple-negative breast cancers

We evaluated TNBC diagnostic core biopsy (pre-treatment) and surgical (post-treatment) specimens from patients treated with neoadjuvant chemotherapy. SOLTI1007 NeoEribulin specimens were obtained from 55 patients who received 4 cycles of neoadjuvant eribulin (55). Comparator specimens were from 15 patients who received standard-of-care neoadjuvant treatment with 4 cycles of adriamycin/cyclophosphamide followed by 4 cycles of paclitaxel (AC-T).

Specimens were multiplex immunostained for 6 EMT markers (Fig. 8A). To enumerate ratios of epithelial and mesenchymal cell types, cells were grouped based on marker expression. In addition, we identified two new phenotypes (KRT8, and ZEB1/VIM) that offer a more comprehensive representation of marker expression patterns. AC-T-treated tumors did not undergo major shifts in subtype proportions. In contrast, eribulin-treated tumors exhibited ∼20% and ∼30% increases in expression of E-Cad/KRT8 and KRT8 epithelial phenotypes, respectively, commensurate with decreases of ∼50% in VIM and ZEB1/VIM mesenchymal phenotypes (Fig. 8B).

**Figure 8:**
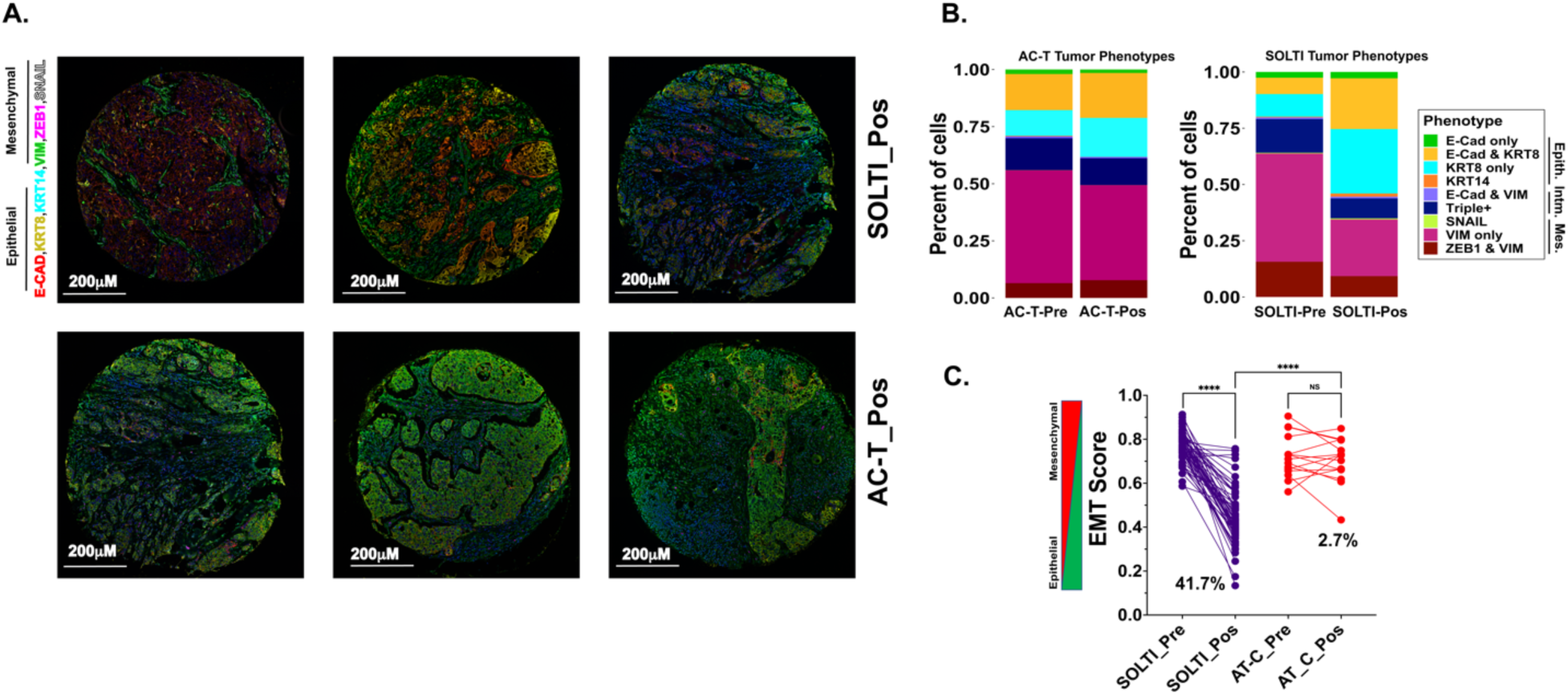
Eribulin reduces EMT in human triple-negative breast cancer. Multiplexed, multi-round TSA staining of EMT markers in triple-negative breast cancer post-treatment with eribulin and AC-T (A) to evaluate distribution of tumor phenotypes by calculating the percent of tumor cells expressing various EMT phenotypes (B) and calculating their EMT scores (C). Tumors were stained using anti-E-CAD (red), anti-KRT8 (yellow), anti-KRT14 (cyan), anti-VIM (green), anti-ZEB1 (magenta), and SNAIL (white) antibodies and counterstained with DAPI for the nuclei (blue). (∗p <0.05, ∗∗p < 0.001, ∗∗∗p > 0.0005, ∗∗∗∗p < 0.0001, Two-Way ANOVA, n=5).

EMT scores were generated for each tumor specimens. Tumors treated with neoadjuvant eribulin showed a 41.7% decrease in EMT score compared to baseline, reflecting as shift towards an epithelial phenotype. In contrast, neoadjuvant AC-T did not significantly alter EMT score (Fig. 8C).

## Discussion

The therapeutic targeting of the EMT has long been pursued as an attractive means of sensitizing tumors that exhibit mesenchymal-like traits to conventional chemotherapy ^2,20–22^. The prevailing understanding of the role of the EMT program in the invasion-metastasis cascade posits that tumors cells that acquire invasive traits by undergoing EMT often endure a reversal of this program via MET at a distant metastatic site, thereby enabling the colonization of foreign tissues by imparting enhanced proliferative traits on tumors cells ^23,24^. Such a notion would argue against the induction of MET as a possible therapeutic avenue for tumors, given the possibility of inadvertently promoting the colonization of tumor cells that have already embarked on the metastatic cascade and disseminated to distant sites prior to therapy. However, our data demonstrating the ability of eribulin to induce regression of established metastases provides evidence that MET induction could prove beneficial in countering metastatic progression. The regression of metastases observed in response to eribulin treatment may be a result of its MET-inducing properties or a combination of this ability and its anti-mitotic effects. The inability of paclitaxel or vinorelbine to eradicate metastases indicates that anti-mitotic effects alone are insufficient to achieve regression of established metastases.

A key question that remains is whether cells of both intermediate/hybrid EMT states and more extreme versions of the EMT are equally susceptible to eribulin-induced MET. The MMTV-PyMT-derived PB3 and MDA-MB-231 cells used in our study have previously been shown to reside in a hybrid/quasi-mesenchymal state ^8,25–27^ with other studies demonstrating the ability to inhibit or reverse the EMT in these cells ^28^. In contrast, the acquisition of epithelial phenotypes upon treatment of more extreme mesenchymal cells such as SUM159 cells with eribulin is more modest (Suppl. Fig. S1F/G), possibly due to their residence in a state that is less permissive to reversion. We have previously observed a similar refractory response to MET induction by activation of PKA signaling in some cell systems ^22^, pointing to a more generalizable inability of cells that have advanced beyond a certain point along the E-M spectrum to revert back and regain epithelial traits. There is ample recent evidence pointing to the aggressive nature of cell residing in the intermediate EMT state, which exhibit enhanced metastatic ability ^16,25,27,29,30^. Thus, whether the therapeutic targeting of the more extreme mesenchymal cell states may ultimately be of any clinical value is still unknown, although recent work suggests that these more mesenchymal cells may retain some plasticity to acquire epithelial traits following treatment with chemotherapy ^30^.

Drug treatment of cell populations typically results in a mixture of outcomes that are influenced in part by the chromatin state of cells. Using a combination of cellular barcoding and gene expression analyses (QISSMET), we have developed a method to study the mode of resistance that drives drug resistance of populations, specifically grouping cells into two categories per prior definitions ^31^ – Darwinian selection, for cells where terminal treatment UMAP state is similar to untreated cells, and Lamarckian induction, for cells where terminal treatment UMAP state is transcriptionally distinct from that of the untreated cells. Our method allows for the uncovering of specific cellular trajectories as they overcome treatment, and it is immediately evident that the same drug can invoke different outcomes in a population of cells, likely determined by the starting chromatin and EMT state of cells. It is interesting to note from the Muller plots in Fig 2H that eribulin treatment imposes a bottleneck that only a small subset of cells can overcome and evolve resistance to by undergoing an shift in their transcriptional and chromatin landscape, reminiscent of previously described cancer stem cell subpopulations that drive cancer relapse following therapy ^20,32,33^.

Through this work, we uncover a novel mechanism that serves to maintain a mesenchymal state in breast cancer cells by altering the transcriptional functionality of the key EMT factor, Zeb1, through its interaction with the Smarcc and Smarcd members of the SWI/SNF chromatin remodeling complex. Several different mechanisms have been previously reported to play a role in regulating Zeb1 repressor function, including chromatin regulators such as CtBP ^34^, LSD1 ^35^, HDAC1/2 ^36^ and BRG1 ^19^. Several of these interactions could constitute attractive therapeutic targets of the EMT program, with previous work demonstrating the utility of HDAC inhibitors such as mocetinostat in reversing Zeb1-associated drug resistance ^37^. While HDAC inhibitors are FDA-approved for certain indications and serve as excellent proof-of-principle epigenetic therapeutics, the lack of a specific protein target has proven to be a hurdle in their success as widely used anti-neoplastic agents. We uncover here that eribulin, a microtubule dynamics inhibitor, could alter the chromatin landscape of breast tumor cells by specifically interfering with the interaction between Zeb1 and core members of the SWI/SNF complex, a potential secondary mechanism of action.

Our studies have significant implications for the development of more effective clinical regimens for breast cancer, specifically those that include neoadjuvant chemotherapy as standard of care. Analysis of tumors from the SOLTI1007 study revealed that treatment of patients with single-agent eribulin in the neoadjuvant setting can induce a reversion of tumor cells to a more epithelial state. We have also observed in our preclinical models that the chromatin-modulating and cellular reprogramming effects of eribulin ensure that pretreatment with this drug sensitizes cells to treatment with subsequent rounds of chemotherapy with other drugs. Thus, there is substantial potential for the use of eribulin pre-treatment as an MET-inducing step, specifically for tumors that exhibit features of increased epithelial-mesenchymal heterogeneity. There are currently no means to stratify patients by the extent of EMT present in their tumor as a biomarker that could inform therapeutic decision-making, with the closest surrogates being an estimate of differentiation state by histopathological examination.

We have recently developed a multi-round multiplexed immunostaining approach to quantify the extent of epithelial-mesenchymal heterogeneity within a given tumor ^16^. Such an approach would enable the identification of patients that may benefit from pretreatment with MET-inducing regimens with drugs such as eribulin prior to receiving standard neoadjuvant chemotherapy. In summary, we have unveiled how a more refined understanding of the mechanism of action of an FDA-approved drug, including its effects on epithelial-mesenchymal plasticity, can open up novel avenues that have the potential to improve clinical outcomes.

## Acknowledgements

We thank the Dartmouth Cancer Center Genomics and Molecular Biology Shared Resource, DartLab Flow Cytometry Shared Resource, Pathology Shared Resource and Mouse Modeling Shared Resource at the Norris Cotton Center. We thank Meredith Brown, Jennifer Fields and Dr. Radu Stan for technical assistance and helpful discussions and Dr. Robert A. Weinberg (Whitehead Institute) for critical reading of the manuscript. Funding and resources for the shared resources were supported in part by a core grant (P30CA023108; Dartmouth Cancer Center). Research reported in this publication was supported through Geisel School of Medicine at Dartmouth’s Center for Quantitative Biology through a grant from the NIGMS Award P20GM130454, an NIH S10 (S10OD025235) grant, a grant from METAvivor (D.R.P), a Prouty Pilot Grant from Friends of the Dartmouth Cancer Center (D.R.P and S.A.G), funding from The Elmer R. Pfefferkorn & Allan U. Munck Education and Research Fund at the Geisel School of Medicine at Dartmouth (D.R.P), and a Sponsored Research Agreement with Eisai Inc. (D.R.P). This work was supported by funding from the NIH R01GM122846 (S.A.G), R00CA201574 (D.R.P.) and R01CA267691 (D.R.P. and T.W.M.).

## Author Contributions

Conception and design: M.B., D.R.P.

**Development of methodology**: M.B., N.B.O., G.A.M., H.L., M.A.M.S., F.W.K., S.H.N., S.A.G., D.R.P.

**Acquisition of data**: M.B., G.A.M., N.B.O., H.L., I.L.C., S.A.G., D.R.P.

**Analysis and interpretation of data**: M.B., N.B.O., G.A.M., M.A.M.S., S.D., H.L., F.W.K., O.M.W., I.L.C., S.H.N., K.E.M., S.A.G., T.W.M., D.R.P.

**Writing, review, and/or editing of the manuscript**: M.B., G.A.M., T.W.M., D.R.P.

**Study supervision**: D.R.P.

## Data and materials availability

The raw and analyzed data were submitted to GEO with accession number GSE207482 and GSE207773. The tutorial and original code are available at: https://github.com/lifebytes/bagheri-et-al_scRNA-ATAC_analysis

## Materials and Methods

### Cell lines and Culture conditions

The MMTV-PyMT derived PB3, MDA-MB-231 and SUM-159 cell lines were gifts from Dr. Bob Weinberg (Whitehead Institute for Biomedical Research). PB3 cells were cultured in DMEM/F12 media supplemented with 5% ABS, L-Glutamine and NEAA. MDA-MB-231 and SUM-159 were cultured in DMEM supplemented with 10% FBS, and F12 supplemented with 10% FBS, respectively, per ATCC guidelines.

### Constructs and plasmids

SMARCD1-Lentiviral (ORFeome) was purchased from GeneCopoeiaTM (EX-A1760-LX304) and used to overexpress SMARCD1 in PB3 and MDA-MB-231 cell lines. shRNAs were designed and synthesized from IDT and cloned into Tet-pLKO-puro vector (Addgene plasmid # 21915) according to the previously used protocols ^38^. CRISPR/Cas9 sgRNAs were designed and synthesized by IDT and cloned into pLenti-CRISPR V2 (Addgene plasmid # 52961) according to the previous used protocols ^39^. To trace metastasis, cells were infected with pHIV-Luc-ZsGreen viral particles (Addgene plasmid # 39196), FACS sorted for ZsGreen expression and used for in vivo experiments. Luciferase *in vivo* imaging was carried out on the IVIS® Lumina III In Vivo Imaging System (PerkinElmer).

Lentiviral particles were produced using standard protocols. Briefly, HEK293T cells were transfected by transfer plasmid and packaging plasmids psPAX2 (addgene #12260) and pMD2.G (addgene #12259) using Lipofectamine 3000 reagent (Thermo Fisher Scientific # L3000015). Viral particles were concentrated using Lenti-X Concentrator (Takara Bio, #631232), and cells were transduced at a high multiplicity of infection with different polybrene concentrations (8ug/mL-30ug/mL).

### Drug Treatment and IC_50_ performance

Stock solutions for eribulin and vinorelbine (10 mM) were prepared using double distilled water, and paclitaxel was prepared using DMSO, further diluted in the base diluent to the appropriate concentrations. The half-maximal inhibitory concentration (IC50) was determined at different concentrations from 50nM to 1500nM and 0.5nM to 100nM using a 2-fold dilution series for PB3 and MDA-MB-231 cells, respectively. Cell viability was assessed using alamarBlue™ Cell Viability Reagent (Thermo Fisher Scientific #DAL1100) and according to manufacturer’s protocols. Briefly, cells were plated in 96-well white clear, flat-bottom microplates (Corning Life Sciences) at a density of 7 × 10^3^ cells per well in 100 µl culture medium supplemented with 10% FBS for 24 hours. Cell viability was assessed 72 hours after drug treatment using 0.2 mg/ml resazurin solutions prepared from resazurin sodium salt (Thermo Fisher Scientific # R12204) dissolved in sterile 1 x PBS (Thermo Fisher Scientific #10010023). The cells were incubated with 10 µl resazurin solution (10% cell culture volume) for four hours at 37 °C. The absorbance was measured with a 560 nm excitation filter and a 590 nm emission filter in a SpectraMax i3x microplate reader (Molecular Devices).

Percentage cell viability was calculated as 100% × (absorbance of treated cells – absorbance of background controls) / (absorbance of matched controls – absorbance of background controls). The IC50 was determined for each compound using the nonlinear regression method (PrismAcademy V9).

### SAβ-Gal, Cell dead and Apoptosis assay

β-galactosidase activity in the PB3 and MDA-MB-231 cells after different treatments were evaluated using the CellEvent™ Senescence Green Flow Cytometry Assay Kit (Thermo Fisher Scientific #C10841) and manufacturer’s protocols, detecting β-galactosidase activity by flow cytometry. Briefly, following different treatments, cells were trypsinized, washed with 1X PBS, resuspended in 100 µL of fixation solution, and incubated for 10 minutes at room temperature. Then cells were washed in 1% BSA in PBS to remove the fixation solution, resuspended in 100 µL of working solution, incubated for 2 hours at 37° C without CO2. Cells were resuspended in 1% BSA in PBS and analyzed on a flow cytometer using a 488-nm laser and 530-nm/30 filter. The mean fluorescence intensity was measured in each experiment.

Cell dead and apoptosis were determined in PB3 and MDA-MB-231 cells using the Dead Cell Apoptosis Kit (Thermo Fisher Scientific #V35114) following different treatments according to the manufacturer’s protocols. Briefly, cells were harvested 72 h after treatment and washed in 1X Annexin binding buffer. Washed cells spun down resuspended in 1X Annexin binding buffer and stained with SYTOX Green, Resazurin, and APC-Annexin V. cells incubated at 37°C in an atmosphere of 5% CO2 for 15 minutes. After the incubation period, 400ul 1X annexin-binding buffer was added to the samples, and data were collected by flow cytometry (Gallios™ Flow Cytometer, Beckman Coulter) at 530 nm and 575 nm using 488 nm excitation, and at 660 nm using 633 nm excitation. Data analysis was carried out on FlowJo V.10.8.1 (BD biosciences).

### Immunoblotting

Protein extraction was performed by pelleting cells and resuspending in cold RIPA buffer (Thermo Fisher Scientific, #89901) supplemented with phosphatase/protease inhibitors (Thermo Fisher Scientific, #78446) and incubating for 1 hour on ice. Then, protein extracts were collected at 20,000 rcf for 10 min in a refrigerated benchtop centrifuge. Protein lysates were quantified using Quick Start™ Bradford Protein Assay kit (BIO-RAD #5000202) and then diluted NuPAGE LDS sample buffer (Life Technologies #NP0007) and NuPAGE™ Sample Reducing Agent (Life Technologies #NP0009) and heated at 72°C before resolving on 4-12% Bis-Tris gradient gels. Gels were either wet or dry transferred using Mini Blot module or iBlot™ 2 Gel Transfer Device, respectively (Thermo Fisher Scientific #B1000 and #IB21001). All primary antibodies were incubated overnight with membranes in TBS blocking buffer supplemented with 0.1% Tween-20 (#), while secondary antibodies (HRP conjugate) were incubated at room temperature with agitation for 1 hour in primary blocking buffer. Membranes were developed using SuperSignal™ Western Blot Substrate (Thermo Fisher Scientific, # A45915). List of antibodies were used provided in the table of materials below.

| Item | Supplier | Cat. Number | Application |
| --- | --- | --- | --- |
| Eribulin | Eisai Inc. |  | Drug treatment of cells |
| Vinorelbine | SAGENT | NDC 25021-204-01 | Drug treatment of cells |
| Paclitaxel | Hospira | NDC 61703-342-09 | Drug treatment of cells |
| APC anti-mouse<br>CD326 (Ep-CAM) | BioLegend | 118214 | Flow cytometry |
| anti-E-cadherin | BD | 610182 | IHF, TSA |
| anti-E-cadherin | CST | 3195 | WB |
| anti-Vimentin | CST | 5741 | WB, TSA, IHF |
| anti-Snail | CST | 3879 | TSA |
| anti-Snail | CST | 3895 | WB |
| anti-Tubulin | CST | 5666 | WB |
| anti-Gapdh | CST | 2118 | WB |
| anti-Zeb1 | Bethyl | A301-922A | WB, IP, CUT&RUN |
| anti-Smarcd1 | Invitrogen | PA5-52049 | WB |
| anti-Smarcc1 | Bethyl | A301-020A | IP |
| anti-Smarcc1 | CST | 11956 | WB |
| anti-Smarcd3 | CST | 62265 | WB |
| anti-H3K4me3 | Invitrogen | MA5-11199 | WB, CUT&RUN |
| anti-H3K27me3 | Invitrogen | MA5-11198 | WB |
| anti-CD104APC | BioLegend | 123612 | Fellow Cytometry |
| anti-CD44PECy7 | BioLegend | 858802 | Fellow Cytometry |
| anti-KRT8 | ThermoFisher | PA5-29607 | TSA, IHF |
| anti-KRT14 | ThermoFisher | MA5-11599 | TSA, IHF |
| Sa-bGal kit | CST | 23833 | Senescent cells<br>detection |
| LIVE/DEAD™<br>Viability/Cytotoxicity<br>Kit | Invitrogen | L3224 | Live-dead and<br>apoptosis detection |
| VivoGlo Luciferin | Promega | P1043 | In vivo imaging |
| Universal Magnetic<br>Co-Ip kit | Active Motif | 54002 | CO-IP |
| CUTANA™<br>CUT&RUN Kit | EpiCypher | 14-1048 | CUT&RUN |
| ATAC-seq kit |  |  | ATAC-seq |
| RNeasy plus kit | Qiagen | 74134 | RNA isolation |
| QIAamp DNA<br>isolation kit | Qiagen | 51304 | DNA isolation |
| MinElute Purification<br>Kit | Qiagen | 28004 | ATAC-seq and<br>CUT&RUN |

### Immunoprecipitation

PB3 and MDA-MB-231 cells were seeded in 150mm cell culture dishes and treated with eribulin for 48 hours for immunoprecipitation. Nuclear fractions were extracted following the manufacturer’s instruction supplied with the Universal Magnetic co-IP kit (54002, Active Motif). 1000μg of proteins were immunoprecipitated using 5.0μg of antibody followed by protein G magnetic beads. The immune complexes were analyzed by western blot.

### Immunofluorescence microscopy

Cells (5×10^3^ cells per chamber) were seeded into 8-chamber (BD Falcon) pretreated culture slides with poly L-lysin (#A3890401, Gibco) and stained according to the manufacturing antibody provider. Briefly, cells were rinsed with ice-cold 1X PBS and fixed with 4% paraformaldehyde or cold acetone-methanol (1:1) for 10 min at room temperature, followed by permeabilization with 0.1% Triton X-100. The cells were subjected to blocking buffer recommended by the manufacturing antibody for 1 hour at room temperature followed by primary antibody incubation overnight at 4 0C. The cells were then washed with cold 1X PBS three times for 3 min each and incubated with Alexa 555-labeled anti-rabbit and Alexa 488-labeled anti-mouse secondary antibody (Thermo Fisher) at room temperature for 1 hour. The cells were washed three times with cold 1X PBS. The cells were examined by fluorescence confocal microscopy (Zeiss LSM 800 with Airyscan (63X)).

### Immunohistofluorescence staining

Before proceeding with the staining protocol, tissue sections from fixed and paraffin-embedded tissues were deparaffinized using xylene. They rehydrated using the decreasing concentration of ethanol (100%, 90%, 70%, and 50% subsequently,) followed by water wash. Antigen retrieval was performed by sodium citrate buffer (pH 6.0) for 20 minutes in a pressure cooker followed by 20 min of cooling at room temperature. Sections were incubated with protein blocking buffer (1% goat serum in 1% BSA) for 1 hour at room temperature. Slides were then incubated at 4°C overnight with primary antibodies. After rinsing the slides in TBST, they were incubated in secondary antibody conjugated to different fluorophores (Alexa 555-labeled anti-rabbit and Alexa 488-labeled anti-mouse secondary antibody (Thermo Fisher)) and DAPI for 1 hour at room temperature. Slides were washed with TBST and mounted using Prolong™ diamond antifade mountant (Thermo Fisher Scientific) and imaged using fluorescence confocal microscopy (Zeiss LSM 800 with Airyscan (63X)).

## In vitro Assays

### Transwell Assay

The invasion rate of different resistant clones was assessed using 24-well Transwell® permeable supports (Costar, Corning Inc., Corning, NY, USA) with polycarbonate (PC) membrane 8.0μm pore size. The transwells were treated by matrigel (BD Bioscience, NJ, USA) for 3 hours and solidified. Cells trypsinized and prepared into 1 × 10^5^ cells/mL dilutions with a serum-free cell culture medium. Then, an appropriate amount of cell suspension was supplemented in the transwell chamber supplied with 0.4ml serum-free media. The lower chamber contained media supplemented with FBS. After incubation for 8 hours, non-invaded cells were removed from the top well using a cotton swab. Cells were fixed using 4% paraformaldehyde and were stained with 0.1% crystal violet. Finally, the invading cells were counted manually.

### Scratch assay

The migration capabilities of different resistant clones were assessed using a scratch wound assay. The cells were seeded into a 6-well tissue culture plate and incubated overnight at 37 °C, at a concentration of 5 × 105 cells per well to nearly confluent cell monolayers. Then, a linear wound was generated in the monolayer with a sterile 100μl plastic pipette tip. Wells were washed with phosphate buffer saline (PBS) to remove debris. Cells were incubated at 37 °C with 5% CO2 and imaged after 2, 4, 8, and 16 hours.

### Generation of PDX and in vivo tumorigenicity

The JAX-98 (TM00098) and JAX-91 (TM00091) models were purchased from The Jackson Laboratory. NCI-140-R (994819) was obtained from the NCI Patient-Derived Models Repository (PDMR). NCCC-470 was obtained from the Dartmouth Cancer Center Mouse Modeling Shared Resource. All PDX models were derived from primary tumor surgical specimens. JAX-98 and NCI-140-R were derived from treatment-naïve patients. NCCC-470 and JAX-91 were derived from patients treated with neoadjuvant chemotherapy.

First, PDXs were expanded primarily by subcutaneous fragment engraftment into 8 to 10 weeks old NOD/SCID mice [NOD.*Cg-Prkdcscid Il2rgtm1Wjl*/SzJ (NSG; Strain #:005557), The Jackson Laboratory]. When tumors reached about 1500 mm^3^, they were harvested and washed with 1X cold PBS containing antibiotics [penicillin (100 U/ml), streptomycin (100 ug/ml), and amphotericin B (0.25 ug/ml)]. Tumors were sliced into 5mm^3^ chunks and cryopreserved or engrafted into the mammary fat pad of NOD/SCID mice for further experiments.

### Tumor Monitoring and Treatment

The growth rate of PDXs were highly variable, Jax-91 and NCI-140 tumors initially appearing in as little as several weeks but NCCC-470 and Jax-98 appeared and grow in a few months. Subcutaneously implanted tumors monitored twice a week and tumor volumes measured with a digital caliper using the formula, Length×Width^2^/2. To have a normalized tumor volume, the tumor volume percentage was calculated using the adapted formula from Gao et al., 2015 (Nature Medicine 21, 1318-1325), V: ((end volume – start volume)/start volume) * 100).

Vinorelbine and paclitaxel were purchased from the Dartmouth-Hitchcock Medical Center pharmacy, and eribulin powder was supplied by Eisai Inc., USA. Eribulin was solubilized in sterile water for injection immediately prior to administration to mice. The solutions were protected from light and were made fresh before injecting each cohort of animals. Vinorelbine (7 mg/kg/bw), paclitaxel (20 mg/kg/bw) and eribulin (1.6 mg/kg/bw) solutions were administered by intraperitoneal injection separately at a dose volume of 200ul twice a week for a total of 6,6, and 4 doses respectively. Tumors were treated when they reached to 500mm^3^. After the first round of treatment one of two engrafted tumors was surgically removed and divided into different sections for downstream analysis. Second round of treatment was started two weeks following surgery with subsequent drug treatment for the same duration as the first. The second tumors were surgically resected, following which mice were held for three months to monitor tumor recurrence and metastasis. Health and body weight were monitored two to three times weekly.

The tumor response rate was determined based on the minimum volume (Vm) and the mean of tumor volume (Va) in each group and categorized based on the following table ^40^.

| **RECIST Category** | **Best Response: Vm** | **Average Response: Va** |
| --- | --- | --- |
| Complete Response (CR) | < -95% | < -40% |
| Partial Response (PR) | < -50% | <-20% |
| Stable Disease (SD) | < 35% | < 30% |
| Progressive Disease (PD) | Anything else | Anything else |

### Proteome Integral Stability Assay

The general PISA procedure ^18^ was adapted as follows: PB3 and MDA-MB 231 cells were cultured as described above in 10cm dishes and treated in triplicate with either DMSO (vehicle control), 300nM eribulin, or 600nM eribulin for 4 hours, followed by harvesting by trypsinization, washing once with PBS and resuspending in PBS to a cell density equivalent to 0.4mg protein/ml PBS. 100mL cell suspension from each replicate of each condition was transferred to 9 PCR tubes in a strip, followed by incubation at different temperatures for 3 minutes in a PCR machine. Samples were removed from the PCR machine and immediately snap-frozen in liquid nitrogen, then thawed at room temperature for 5 minutes in a water bath. This process was repeated twice, followed by centrifugation at 21,000 x g at 4C for 10 minutes. 50ul of supernatant from each temperature condition was combined in a clean Eppendorf tube, followed by the addition of 1ml 9M urea/50mM Tris pH 8.1/150mM NaCl/2mM DTT, heating at 45C for 15 minutes to reduce each sample and alkylation with 7mM iodoacetamide for 1 hour in the dark. The alkylation reaction was quenched with an additional 2mM DTT for 15 minutes, followed by dilution in 1.5ml 50mM Tris pH 8.1/150mM NaCl and digestion with 1mg sequencing-grade trypsin overnight at 30C. Each sample was desalted on an OASIS mHLB plate (2mg sorbent) and quantified by peptide BCA assay (Thermo) per manufacturer’s directions. The equivalent of 40mg peptide from each replicate of each condition was labeled using unique isobaric labeling tags (TMT 10-plex reagents, ThermoFisher Scientific) by resuspension in 30ug 166mM EPPS, pH 9, and addition of TMT reagent (80ug reagent in 8ul dry acetonitrile) for 1 hour, followed by quenching with 1% hydroxylamine for 20 minutes at room temperature. Each multiplex sample was combined in a single Eppendorf tube, followed by the addition of 600ml 1% TFA and desalting on an OASIS C18 plate (10mg sorbent). The desalted multiplex was dried by vacuum centrifugation and separated on a pentafluorophenyl (PFP) reverse-phase column (Waters) into 48 distinct fractions ^41^, concatenated into 16 individual combined fractions by sequential addition, dried by vacuum centrifugation, and analyzed by LC-MS/MS on a Proxeon Easy-nLC 1200-Orbitrap Fusion Lumos mass spectrometer platform using SPS-MS3-based quantification. The resulting tandem mass spectra were data-searched using the Comet algorithm (56) against the human database (UniProt; www.uniprot.org), filtered to a 1% false discovery rate using the target-decoy strategy ^42^, and reported. Quantification was accomplished using in-house software to extract reporter ion intensities per-peptide and summed across all peptides for each unique UniProt identifier. Each channel was normalized across all proteins to the total intensity of the least abundant channel and proteins were adjusted accordingly. Final protein stability ratios were reported as delta-Sm values precisely described ^18^.

### RNA-seq data Library Preparation

The cells were washed twice and harvested in 1X PBS, and total RNA was extracted using RNeasy Plus Mini Kit (QIAGEN #74136). The purified RNA was used for reverse transcription with High-Capacity cDNA Reverse Transcription Kits (Applied Biosystems, #4374967). Briefly, 1.5 ug RNA sample was applied in a 20 uL reaction and incubated at 25 C for 10 min, followed by 120 min at 37 C. Then, the reverse transcriptase was inactivated by heating to 85 C for 5 min. Finally, the cDNA product was used for next-generation sequencing (NGS) library preparation. RNA-seq data were analyzed as previously described ^16^.

### Pre-processing of ATAC-seq data

Methodology adapted from (1). In short, cells were washed with cold 1X PBS and trypsinized. After the cells were counted 100,000 cells were resuspended in cold 1X PBS and centrifuged at 500*g* for 5min. Cells were resuspended in 1 ml of cold ATAC-seq resuspension buffer (RSB, 10 mM Tris-HCl pH 7.4, 10 mM NaCl, and 3 mM MgCl_2_ in water) and centrifuged at 500*g* for 5min in a precooled fixed angel centrifuge. Nuclei were isolated by resuspending cells in 100μl of ATAC-seq lysis buffer (RSB containing 0.1% NP40, 0.1% Tween-20, and 0.01% digitonin) by pipetting up and down 5 times. Cells suspension were incubated 3 min on ice and washed with 1ml of ATAC-seq RSB containing 0.1% Tween-20 (without NP40 or digitonin). Nuclei spun for 10 min at 500g in a pre-cooled centrifuge, the supernatant was removed carefully, and nuclei were resuspended in 100μl of transposition mix (50μl 2× TD buffer, 5μl transposase (100 nM final), 33μl PBS, 1μl 1% digitonin, 1μl 10% Tween-20, and 10μl water). The transposition reactions were incubated at 37 °C for 30 min in a thermomixer with shaking at 1,000 r.p.m. RNA-seq data was analyzed as previously described ^16^.

### Pre-processing of CUT&RUN data

CUT&RUN was performed using CUTANA™ Kit (EpiCypher #14-1048) according to the manufacturer’s recommendation with minor modifications. Briefly, 5×105 cells were washed and bound to concanavalin A-coated magnetic beads. The cells were then permeabilized with Wash Buffer (20 mM HEPES pH 7.5, 150 mM NaCl, 0.5 mM spermidine, and 1x Roche Complete Protease Inhibitor, EDTA-free) containing 0.025% digitonin (Digitonin Buffer) and 2 mM EDTA and incubated with primary antibody (anti-ZB1, anti-H3K4me3 or IgG isotype control) overnight at 4°C. The cell-bead slurry was washed twice with Digitonin Buffer and incubated with 1x Protein-A/G-MNase (pAG-MNase) in Digitonin Buffer for 10 minutes at room temperature. The slurry was washed twice with Digitonin Buffer and incubated in Digitonin Buffer containing 2 mM CaCl2 for 2 hours at 4°C to activate pAG-MNase digestion. The digestion was stopped by addition of 2x Stop Buffer (340 mM NaCl, 20 mM EDTA, 4 mM EGTA, 50 µg/mL RNase A, 50 µg/mL glycogen), and the sample was incubated for 10 minutes at 37°C to release chromatin to the supernatant and degrade RNA. The supernatant was recovered, and DNA was isolated using MinElute Reaction Cleanup Kit (Qiagen # 28206). Isolated CUT&RUN DNA fragments were quantified by Qubit and 5-10ng used for library preparation with the NEB Ultra II DNA Kit. Library amplification was performed using the modified PCR cycling conditions described in Step 39 of the Epicypher CUTANA^TM^ (EpiCypher) protocol. Libraries individually barcoded and pooled for sequencing on a NextSeq500 Mid Output flow cell to generate 10M, 50bp paired-end reads per sample.

### RNA-seq, ATAC-seq, & CUT&RUN data analysis

Raw sequence quality of reads was determined using FastQC (v0.11.9). Reads were trimmed using Cutadapt (v2.4) for adapter sequences with parameters “--nextseq-trim 20 ––max-n 0.8 –– trim-n-m 1”. Reads were mapped to hg38 (for human samples) or mm10 (for mouse samples) using Bowtie2 (v2.4.2) ^43^ with parameters “--local ––no-mixed –-no-discordant”. Alignments coordinate sorted with samtools (v1.11) ^44^, and unmapped or multi-mapping reads were filtered using sambamba (v0.8.0) ^45^. MarkDuplicates (Picard Tools) was used to identify and remove duplicate reads. Normalized signal tracks were generated using deepTools (v 3.3.0) command BamCoverage with parameters “--binSize 20 ––smoothLength 60 –-normalizeUsing CPM –– centerReads ––extendReads”. Command “bamPEFragmentSize” (deepTools) was used to confirm expected fragment length distributions were present in each sample, and plotFingerprint was used to confirm genomic enrichment in IP samples over corresponding IP controls (IgG). Peaks were called using the MACS2 (v2.2.7.1) call peak command in narrowpeak mode with parameters “-f BAMPE ––keep-dup all –q 0.05”. Corresponding IgG IP controls were specified using option –c. Effective genome size was provided using option –g, with a value of 2913022398 for human data, and 2652783500 for mouse. IP signal was assessed using Fraction of reads in peaks (FRiP), calculated for each sample. Called peaks were filtered against the ENCODE human blacklist (human: ENCFF356LFX, mouse: ENCFF547MET) and further restricted to peaks demonstrating a 2-fold or greater signal increase relative to control (IgG) samples. Reproducible peak sets were identified for replicate pairs using BEDTools (v 2.30.0) command “intersect” with parameter “-a”. Peak sets were annotated using the *annotatePeak()* function (ChIPseeker R-package). Promoters were defined as transcriptional start site (TSS) +/-1kb. R-packages TxDb.Hsapiens. UCSC.hg38. knownGene and TxDb.Mmusculus. UCSC.mm10. knownGene were used to define genomic features for human and mouse peaks, respectively. Any peak > 10kb from the closest gene were annotated as “intergenic” peaks. A consensus peak set was generated by merging overlapping peaks called across all samples. Read counts in each consensus peak region were counted from BAM files using the featureCounts() function (Rsubread package) with options “isPairedEnd=TRUE, countMultiMappingReads=FALSE”. DESeq2 ^46^ was used to perform differential binding analysis on the matrix of read counts from consensus peak regions. Adequate dispersion trends were confirmed during model fitting for all analyses. A Benjamini-Hochberg adjusted *P*-value threshold of < 0.05 (Wald-test) was used to denote statistically significant peaks.

### Multiplexed TSA staining

Methodology adapted from (15). Briefly, four different PDXs, the two naïve (NCI-140 and JAX-91) and two pretreated (NCCC-470 and JAX-98), were selected and treated with different chemotherapeutics drugs (paclitaxel and eribulin) as a single treatment or sequential double treatment regimens. Five cores were selected and stained from untreated and treated samples with a panel of canonical EMT markers. Human specimens of triple-negative breast cancer, both pre-and post-treatment, were obtained from two sources: the SOLTI-1007-NeoEribulin cohort study (n = 55) and the Dartmouth Cancer Center pathology shared resources (n = 15). The core needle biopsy samples that were collected pre-treatment were used directly for TSA staining. Post-treatment samples were processed into tissue microarrays (TMA) with five cores per sample and were then stained with a panel of epithelial-to-mesenchymal transition (EMT) markers, including anti-Snail (CST #3895, dilution 1:400), anti-KRT8 (Invitrogen PA5-29607, dilution 1:300), anti-KRT14 (Invitrogen MA5-11599, dilution 1:1000), anti-Vimentin (CST #5741, dilution 1:500), anti-E-cadherin (BD #610182, dilution 1:500), and anti-ZEB1 (Invitrogen PA5-82982, dilution 1:1000) antibodies. The PerkinElmer OPAL Assay Development Guide (August 2017) adopted the optimization of the antibodies and multiplexed staining procedure.

### Image Analysis

Methodology adapted from ^16^. Slides were scanned using 20x objectives with the PerkinElmer Vectra 3 automated multispectral imaging system. Images were imported into InForm analysis software (PerkinElmer), and analysis was performed according to manufacturer instructions and methods adapted from ^16^. Cells were phenotyped based on one or multiple markers (E-cadherin only, KRT8 & E-cadherin, KRT14 only, Triple positive (KRT8+E-cadherin+ Vimentin), Snail only, Vimentin only, and Vimentin+ZEB1) and validated by marker distribution. Entire Cell Mean Fluorescent units were extracted for each marker and normalized as a percentile of maximum and minimum fluorescence across all cells in all images.

### Heterogeneity and EMT Scores

Recently we developed a pipeline to analyze data generated from multiplexed TSA staining. This study used the same pipeline to evaluate tumor heterogeneity and EMT distribution among different PDXs and treatments ^16^.

## Cell barcoding and analysis

### Lentiviral barcoding and selection of successfully barcoded cells

PB3 cells were grown to an estimated 70% confluency, and trypsinized to obtain a single cell suspension. The cells were counted, and 200 000 cells were plated in a 6cm tissue culture dish, along with 6ug/ml Polybrene transfection reagent (Millipore, Cat #TR-1003-G, Lot #2129807), and were incubated at 37°C for one hour. A lentiviral barcode library (CloneTracker XPTM 10M Barcode-3’ Library with RFP-puro (Packaged), Cellecta #BCXP10M3RP-V), was then added at an MOI of 0.5, and the cells and virus were further incubated at 37°C overnight. After 24 hours, the media was replaced with fresh media, and the cells were allowed to grow for an additional two days. The cells were then trypsinized to obtain a single cell suspension, and flow sorted using the FACSAria III cell sorter to select for RFP expressing, successfully barcoded cells. The collected cells were then re-plated and allowed to grow until the plate reaches confluency. Any cells not used for further experiments were frozen down for long term storage.

### Starting a founder population of barcoded cells

Since the 10x genomics platform could only run a maximum of 10 000 cells per sample, a founder population of at most 10 000 barcodes was first obtained to ensure consistency of barcode distribution across samples. This was done by counting 10 000 cells and plating them in one well of a 24 well plate. This founder population was allowed to grow by scaling up the plate size until a sufficient number of cells are obtained to carry out the drug treatment experiments.

### Drug treatment of barcoded cells

Barcoded PB3 cells were counted, and 80 000 cells were seeded into each well of a 12 well plate. The cells were incubated at 37°C overnight, and 500nM eribulin or 100nM paclitaxel were added. Cells were incubated with the drug for 72 hours, before being replaced with fresh, drug free media. The cells were allowed to recover for 48 hours. Before the next round of treatment, the cells were re-plated into 6 well plates to allow for more space for the cells to grow. As the paclitaxel treated cells seemed to be resistant to 100nM of the drug, the dosage was increased to 250nM for subsequent treatment rounds. The drug treatment was repeated for an additional two rounds, with a portion of cells frozen down after each treatment to represent the different treatment cycles. To drive the eribulin-induced epithelial transition, the eribulin-treated cells were subjected to one additional round of treatment.

### Single-cell omics

Cell cultures containing treated or untreated cells were trypsinized, washed in 1x PBS + 0.04% BSA and counted on a Cellometer K2 instrument (Nexcellom Bioscience). Single cell suspensions were processed using the 10x Genomics NextGem Single Cell RNA-seq v3.1 and Single Cell ATAC-seq v1, with or without multiplexing using lipid-oligo hashtags (Table X), following manufacturer instructions for nuclei isolation (scATAC) library preparation and sequencing. A second batch of untreated and Eri4 samples were run on the 10x Genomics Multiome ATAC + Gene Expression assay. For retrieval of cell barcodes, 10ul of amplified cDNA was subjected to 10 cycles of PCR using the 10x Genomics SI-PCR primer (5’AATGATACGGCGACCACCGAGATCTACACTCTTTCCCTACACGACGC*T*C) and BC_RPI_X_Fwd (CAAGCAGAAGACGGCATACGAGATNNNNNNGTCTCGTGGGCTCGGAGATGTGTATAAGAGACAGACCGAACGCAACGCACGCA), where ‘N’ denotes sample-specific index sequences. Amplicons were purified with SPRI Select beads (Beckman Coulter), pooled with scRNAseq libraries and sequenced to a target depth of 5,000 reads/cell.

### Barcode whitelist preparation

The lentiviral barcode library was purchased from Cellecta (CloneTracker XPTM 10M Barcode-3’ Library with RFP-puro (Packaged), Cellecta #BCXP10M3RP-V). We have prepared a whitelist of barcode by combining BC14 and BC30 (BC14sequence+BC30sequence) barcodes. In custom barcode only sequencing, barcodes are expected in the following format, Custom barcode = 19bp constant + BC14 sequence + TGGT + BC30 sequence We have prepared these barcodes in various formats as per the requirement of downstream tools using in-house scripts. These reference files are used for feature barcode demultiplexing in downstream for custom barcodes FASTQ files.

#### Single cell/nuclei transcriptome data analysis

Eight libraries for single cell/nuclei transcriptome data were obtained in 3 batches namely Run1, Run2 and Run3. Each batch contains different treatment and dosage samples sequenced as listed in supplementary table S1 with their summary statistics. While Run1 and Run2 data were from single cell 3’ V3 sequencing (scRNA) protocol, Run3 was single nuclei RNA+ATAC V1 sequencing (snRNA multiome) protocol. Hence, FASTQ and count matrix were generated using 10x Genomics cellranger-3.1.0, 10x Genomics cellranger-4.0.0 ^47^ and 10x Genomics cellranger-arc-1.0.0 ^48^ for Run1, Run2 and Run3 respectively. Cellranger reference mm10 was used as reference for all runs. For Run1 and Run2, whitelist custom barcodes (as detailed in previous section) were used in cellranger “--feature-ref” parameter for custom barcodes FASTQ. As cellranger-arc pipeline doesn’t support “--feature-ref” barcoding, custom barcode FASTQ files were processed using CITE-seq-Count v1.4.3 tool (Roelli et al., 2019).

Count matrices were analysed using Seurat v3.1.4 pipeline ^49^. Briefly, low quality cells (number of genes expressed < 200, percentage of UMIs in mitochondrial and ribosomal genes individually > 50) are filtered out. Using selected variably expressing genes and significant PCs from PCA, clustering and UMAP projection were generated. Further, all the samples are merged into one object to illustrate the differences due to treated drug and their varying dosage effect. Post merging, batch effects due to run and technology differences were regressed out. Pseudotime analysis was done against all samples merged dataset using Monocle3 v0.2.1 ^11–13^ while considering “Untreated” cells as the root cells. To represent the relative frequency of each barcode over pseudotime, Muller plot representation was used ^50^. Relative frequencies were calculated as the ratio between number of cells associated with a barcode at a given pseudotime (binned) and the sum of all cells (from all barcodes) at the same pseudotime bin. Muller plot is accompanied with line chart illustrating the total number of cells at any given pseudotime.

### Single cell/nuclei ATAC data analysis

Four libraries for single cell/nuclei ATAC data were obtained in 2 batches namely Run1 and Run3 using scATAC V1 and snATAC (multiome) V1 protocol respectively. Sample and summary statistics are provided in supplementary table S2. FASTQ and peak count matrix were generated using 10x Genomics cellranger-atac-1.2.0 and 10x Genomics cellranger-arc-1.0.0 for Run1 and Run3 respectively. Cellranger reference mm10 was used as reference for both the runs. Custom barcode FASTQ files were processed using CITE-seq-Count v1.4.3 tool.

Peak count matrices were analysed using Signac v1.1.1 pipeline. Briefly, low quality cells (fragments in peak regions < 1000 and >75000, percentage of UMIs in peaks > 20, ratio of UMIs in blacklist regions to that of peak regions > 0.05, nucleosome signal > 10 and TSS enrichment < 2) are filtered out. Using selected top features and 2 to 30 PCs from LSI, clustering and UMAP projection were generated. Further, all the samples are integrated into one object using Harmony R package ^51^ to illustrate the differences due to treated drug and their varying dosage effect. In harmony, batch effects due to run and technology differences were regressed out.

### Integration and Custom barcode classification

To assign each cell to a specific custom barcode, we have applied MULTIseqDemux method ^52^ from Seurat pipeline. Though the method has managed to differentiate the barcodes, this method was originally designed for multiple samples demultiplexing, where background estimation is appropriated for each cell from all the barcodes. Here, the enrichment level for each barcode can vary, and it is appropriate to calculate the background for each barcode from all the cells. We deployed an in-house developed R script using binomial distribution, that estimates background for each barcode from all the cells. We have considered only doublets and negatives annotated cells from MULTIseqDemux annotation. Using this approach, we were able to re-annotate and increase the number of singlets. Finally, RNA and ATAC datasets were integrated with TransferData module in Seurat pipeline.

#### Selection vs induction identification

To identify and illustrate the cells that undergo selection (Treated and Untreated cells clustering together) or induction (Treated and Untreated cells clustering far apart) process upon treatment, all cells in each barcode were analysed using Jaccard index and Euclidean distance-based methods. Euclidean distance is calculated between median points of cells from a sample per barcode in their UMAP space. For differences within samples, we have used Jaccard index method from scclusteval v1.0 ^53^, where cells of each barcode were re-clustered using density clustering method (with same variable genes from original integrated dataset). Based on the number of cells for each sample (treated or untreated) in various clusters, the Jaccard index is calculated. In density clustering, proximal cells are assigned to one cluster. Ideally, if all the cells are similar in a given scenario, then they will cluster closely and be assigned into one cluster.

### Statistical analysis

Data sets were analyzed by unpaired *t*-test or multiple comparisons one-way ANOVA or two-way ANOVA according to the experiment using GraphPad Prism software. ∗p <0.05, ∗∗p < 0.001, ∗∗∗p > 0.0005, ∗∗∗∗p < 0.0001.

### Study approval

This study was carried out in accordance with the recommendations in the Guide for the Care and Use of Laboratory Animals from the National Institutes of Health (NIH) IACUC. The protocol was approved by the IACUC at NCCC (protocol # 0002119). Mice were euthanized when they became moribund or when they reached defined study end points. All relevant ethical regulations were followed as approved by the Dartmouth Cancer Center.

## References

1 Nieto, M. A., Huang, R. Y., Jackson, R. A. & Thiery, J. P. Emt: 2016. Cell 166, 21–45, doi:10.1016/j.cell.2016.06.028 (2016).

2 Shibue, T. & Weinberg, R. A. EMT, CSCs, and drug resistance: the mechanistic link and clinical implications. Nat Rev Clin Oncol 14, 611–629, doi:10.1038/nrclinonc.2017.44 (2017).

3 Cortes, J. et al. Eribulin monotherapy versus treatment of physician’s choice in patients with metastatic breast cancer (EMBRACE): a phase 3 open-label randomised study. Lancet 377, 914–923, doi:10.1016/S0140-6736(11)60070-6 (2011).

4 Smith, J. A. et al. Eribulin binds at microtubule ends to a single site on tubulin to suppress dynamic instability. Biochemistry 49, 1331–1337, doi:10.1021/bi901810u (2010).

5 Yoshida, T. et al. Eribulin mesilate suppresses experimental metastasis of breast cancer cells by reversing phenotype from epithelial-mesenchymal transition (EMT) to mesenchymal-epithelial transition (MET) states. Br J Cancer 110, 1497–1505, doi:10.1038/bjc.2014.80 (2014).

6 Dybdal-Hargreaves, N. F., Risinger, A. L. & Mooberry, S. L. Regulation of E-cadherin localization by microtubule targeting agents: rapid promotion of cortical E-cadherin through p130Cas/Src inhibition by eribulin. Oncotarget 9, 5545–5561, doi:10.18632/oncotarget.23798 (2018).

7 Kaul, R., Risinger, A. L. & Mooberry, S. L. Eribulin rapidly inhibits TGF-beta-induced Snail expression and can induce Slug expression in a Smad4-dependent manner. Br J Cancer 121, 611–621, doi:10.1038/s41416-019-0556-9 (2019).

8 Dongre, A. et al. Epithelial-to-Mesenchymal Transition Contributes to Immunosuppression in Breast Carcinomas. Cancer Res 77, 3982–3989, doi:10.1158/0008-5472.CAN-16-3292 (2017).

9 Chang, M. T. et al. Identifying transcriptional programs underlying cancer drug response with TraCe-seq. Nat Biotechnol 40, 86–93, doi:10.1038/s41587-021-01005-3 (2022).

10 Emert, B. L. et al. Variability within rare cell states enables multiple paths toward drug resistance. Nat Biotechnol 39, 865–876, doi:10.1038/s41587-021-00837-3 (2021).

11 Cao, J. et al. The single-cell transcriptional landscape of mammalian organogenesis. Nature 566, 496–502, doi:10.1038/s41586-019-0969-x (2019).

12 Qiu, X. et al. Reversed graph embedding resolves complex single-cell trajectories. Nat Methods 14, 979–982, doi:10.1038/nmeth.4402 (2017).

13 Trapnell, C. et al. The dynamics and regulators of cell fate decisions are revealed by pseudotemporal ordering of single cells. Nat Biotechnol 32, 381–386, doi:10.1038/nbt.2859 (2014).

14 Husemann, Y. et al. Systemic spread is an early step in breast cancer. Cancer Cell 13, 58–68, doi:10.1016/j.ccr.2007.12.003 (2008).

15 Rhim, A. D. et al. EMT and dissemination precede pancreatic tumor formation. Cell 148, 349–361, doi:10.1016/j.cell.2011.11.025 (2012).

16 Brown, M. S. et al. Dynamic plasticity within the EMT spectrum, rather than static mesenchymal traits, drives tumor heterogeneity and metastatic progression of breast cancers. bioRxiv, 2021.2003.2017.434993, doi:10.1101/2021.03.17.434993 (2021).

17 Berest, I. et al. Quantification of Differential Transcription Factor Activity and Multiomics-Based Classification into Activators and Repressors: diffTF. Cell Rep 29, 3147–3159 e3112, doi:10.1016/j.celrep.2019.10.106 (2019).

18 Gaetani, M. et al. Proteome Integral Solubility Alteration: A High-Throughput Proteomics Assay for Target Deconvolution. J Proteome Res 18, 4027–4037, doi:10.1021/acs.jproteome.9b00500 (2019).

19 Sanchez-Tillo, E. et al. ZEB1 represses E-cadherin and induces an EMT by recruiting the SWI/SNF chromatin-remodeling protein BRG1. Oncogene 29, 3490–3500, doi:10.1038/onc.2010.102 (2010).

20 Gupta, P. B. et al. Identification of selective inhibitors of cancer stem cells by high-throughput screening. Cell 138, 645–659, doi:10.1016/j.cell.2009.06.034 (2009).

21 Ognjenovic, N. B. et al. Limiting Self-Renewal of the Basal Compartment by PKA Activation Induces Differentiation and Alters the Evolution of Mammary Tumors. Dev Cell 55, 544–557 e546, doi:10.1016/j.devcel.2020.10.004 (2020).

22 Pattabiraman, D. R. et al. Activation of PKA leads to mesenchymal-to-epithelial transition and loss of tumor-initiating ability. Science 351, aad3680, doi:10.1126/science.aad3680 (2016).

23 Ocana, O. H. et al. Metastatic colonization requires the repression of the epithelial-mesenchymal transition inducer Prrx1. Cancer Cell 22, 709–724, doi:10.1016/j.ccr.2012.10.012 (2012).

24 Tsai, J. H., Donaher, J. L., Murphy, D. A., Chau, S. & Yang, J. Spatiotemporal regulation of epithelial-mesenchymal transition is essential for squamous cell carcinoma metastasis. Cancer Cell 22, 725–736, doi:10.1016/j.ccr.2012.09.022 (2012).

25 Bierie, B. et al. Integrin-beta4 identifies cancer stem cell-enriched populations of partially mesenchymal carcinoma cells. Proc Natl Acad Sci U S A 114, E2337–E2346, doi:10.1073/pnas.1618298114 (2017).

26 Dongre, A. et al. Direct and Indirect Regulators of Epithelial-Mesenchymal Transition-Mediated Immunosuppression in Breast Carcinomas. Cancer Discov 11, 1286–1305, doi:10.1158/2159-8290.CD-20-0603 (2021).

27 Pastushenko, I. et al. Identification of the tumour transition states occurring during EMT. Nature 556, 463–468, doi:10.1038/s41586-018-0040-3 (2018).

28 Vijay, G. V. et al. GSK3beta regulates epithelial-mesenchymal transition and cancer stem cell properties in triple-negative breast cancer. Breast Cancer Res 21, 37, doi:10.1186/s13058-019-1125-0 (2019).

29 Jolly, M. K. et al. Implications of the Hybrid Epithelial/Mesenchymal Phenotype in Metastasis. Front Oncol 5, 155, doi:10.3389/fonc.2015.00155 (2015).

30 Luond, F. et al. Distinct contributions of partial and full EMT to breast cancer malignancy. Dev Cell 56, 3203–3221 e3211, doi:10.1016/j.devcel.2021.11.006 (2021).

31 Boumahdi, S. & de Sauvage, F. J. The great escape: tumour cell plasticity in resistance to targeted therapy. Nat Rev Drug Discov 19, 39–56, doi:10.1038/s41573-019-0044-1 (2020).

32 Chen, J. et al. A restricted cell population propagates glioblastoma growth after chemotherapy. Nature 488, 522–526, doi:10.1038/nature11287 (2012).

33 Diehn, M. et al. Association of reactive oxygen species levels and radioresistance in cancer stem cells. Nature 458, 780–783, doi:10.1038/nature07733 (2009).

34 Postigo, A. A. & Dean, D. C. ZEB represses transcription through interaction with the corepressor CtBP. Proc Natl Acad Sci U S A 96, 6683–6688, doi:10.1073/pnas.96.12.6683 (1999).

35 Wang, J. et al. Opposing LSD1 complexes function in developmental gene activation and repression programmes. Nature 446, 882–887, doi:10.1038/nature05671 (2007).

36 Aghdassi, A. et al. Recruitment of histone deacetylases HDAC1 and HDAC2 by the transcriptional repressor ZEB1 downregulates E-cadherin expression in pancreatic cancer. Gut 61, 439–448, doi:10.1136/gutjnl-2011-300060 (2012).

37 Meidhof, S. et al. ZEB1-associated drug resistance in cancer cells is reversed by the class I HDAC inhibitor mocetinostat. EMBO Mol Med 7, 831–847, doi:10.15252/emmm.201404396 (2015).

38 Wiederschain, D. et al. Single-vector inducible lentiviral RNAi system for oncology target validation. Cell Cycle 8, 498–504, doi:10.4161/cc.8.3.7701 (2009).

39 Shalem, O. et al. Genome-scale CRISPR-Cas9 knockout screening in human cells. Science 343, 84–87, doi:10.1126/science.1247005 (2014).

40 Eisenhauer, E. A. et al. New response evaluation criteria in solid tumours: revised RECIST guideline (version 1.1). Eur J Cancer 45, 228–247, doi:10.1016/j.ejca.2008.10.026 (2009).

41 Grassetti, A. V., Hards, R. & Gerber, S. A. Offline pentafluorophenyl (PFP)-RP prefractionation as an alternative to high-pH RP for comprehensive LC-MS/MS proteomics and phosphoproteomics. Anal Bioanal Chem 409, 4615–4625, doi:10.1007/s00216-017-0407-6 (2017).

42 Elias, J. E. & Gygi, S. P. Target-decoy search strategy for increased confidence in large-scale protein identifications by mass spectrometry. Nat Methods 4, 207–214, doi:10.1038/nmeth1019 (2007).

43 Langmead, B. & Salzberg, S. L. Fast gapped-read alignment with Bowtie 2. Nat Methods 9, 357–359, doi:10.1038/nmeth.1923 (2012).

44 Li, H. et al. The Sequence Alignment/Map format and SAMtools. Bioinformatics 25, 2078–2079, doi:10.1093/bioinformatics/btp352 (2009).

45 Tarasov, A., Vilella, A. J., Cuppen, E., Nijman, I. J. & Prins, P. Sambamba: fast processing of NGS alignment formats. Bioinformatics 31, 2032–2034, doi:10.1093/bioinformatics/btv098 (2015).

46 Love, M. I., Huber, W. & Anders, S. Moderated estimation of fold change and dispersion for RNA-seq data with DESeq2. Genome Biol 15, 550, doi:10.1186/s13059-014-0550-8 (2014).

47 Zheng, G. X. et al. Massively parallel digital transcriptional profiling of single cells. Nat Commun 8, 14049, doi:10.1038/ncomms14049 (2017).

48 Satpathy, A. T. et al. Massively parallel single-cell chromatin landscapes of human immune cell development and intratumoral T cell exhaustion. Nat Biotechnol 37, 925–936, doi:10.1038/s41587-019-0206-z (2019).

49 Stuart, T. et al. Comprehensive Integration of Single-Cell Data. Cell 177, 1888–1902 e1821, doi:10.1016/j.cell.2019.05.031 (2019).

50 Farahpour, F., Saeedghalati, M. & Hoffmann, D. MullerPlot: Generates Muller Plot from Population/Abundance/Frequency Dynamics Data. https://cran.r-project.org/package=MullerPlot (2016).

51 Korsunsky, I. et al. Fast, sensitive and accurate integration of single-cell data with Harmony. Nat Methods 16, 1289–1296, doi:10.1038/s41592-019-0619-0 (2019).

52 McGinnis, C. S. et al. MULTI-seq: sample multiplexing for single-cell RNA sequencing using lipid-tagged indices. Nat Methods 16, 619–626, doi:10.1038/s41592-019-0433-8 (2019).

53 Tang, M. et al. Evaluating single-cell cluster stability using the Jaccard similarity index. Bioinformatics 37, 2212–2214, doi:10.1093/bioinformatics/btaa956 (2021).

54 Pascual T, Oliveira M, Villagrasa P, Ortega V, Paré L, Bermejo B, Morales S., et al. Neoadjuvant eribulin in HER2-negative early-stage breast cancer (SOLTI-1007-NeoEribulin): a multicenter, two-cohort, non-randomized phase II trial. NPJ Breast Cancer 2021;25;7(1):145. doi:10.1038/s41523-021-00351-4.

55 Eng JK, Jahan TA, Hoopmann MR. Comet: an open-source MS/MS sequence database search tool. Proteomics 2013;13(1):22–4. doi:10.1002/pmic.201200439.

