## Supplementary FIgures and Legends for "Pharmacological Induction of mesenchymal-epithelial transition chemosensitizes breast cancer cells and prevents metastatic progression"

Supplemental Information

### Therapeutic induction of mesenchymal-epithelial transition via epigenetic reprogramming curtails metastatic progression and sensitizes breast cancers to treatment

Meisam Bagheri, Gadisti Aisha Mohamed, Mohammed Ashick Mohamed Saleem, Nevena B. Ognjenovic, Hanxu Lu, Fred W. Kolling 4th, Owen M. Wilkins, Subhadeep Das, Ian S. La Croix, Shivashankar H. Nagaraj, Kristen E. Muller, Scott A. Gerber, Todd W. Miller, Diwakar R. Pattabiraman

**Supplementary Fig 1**

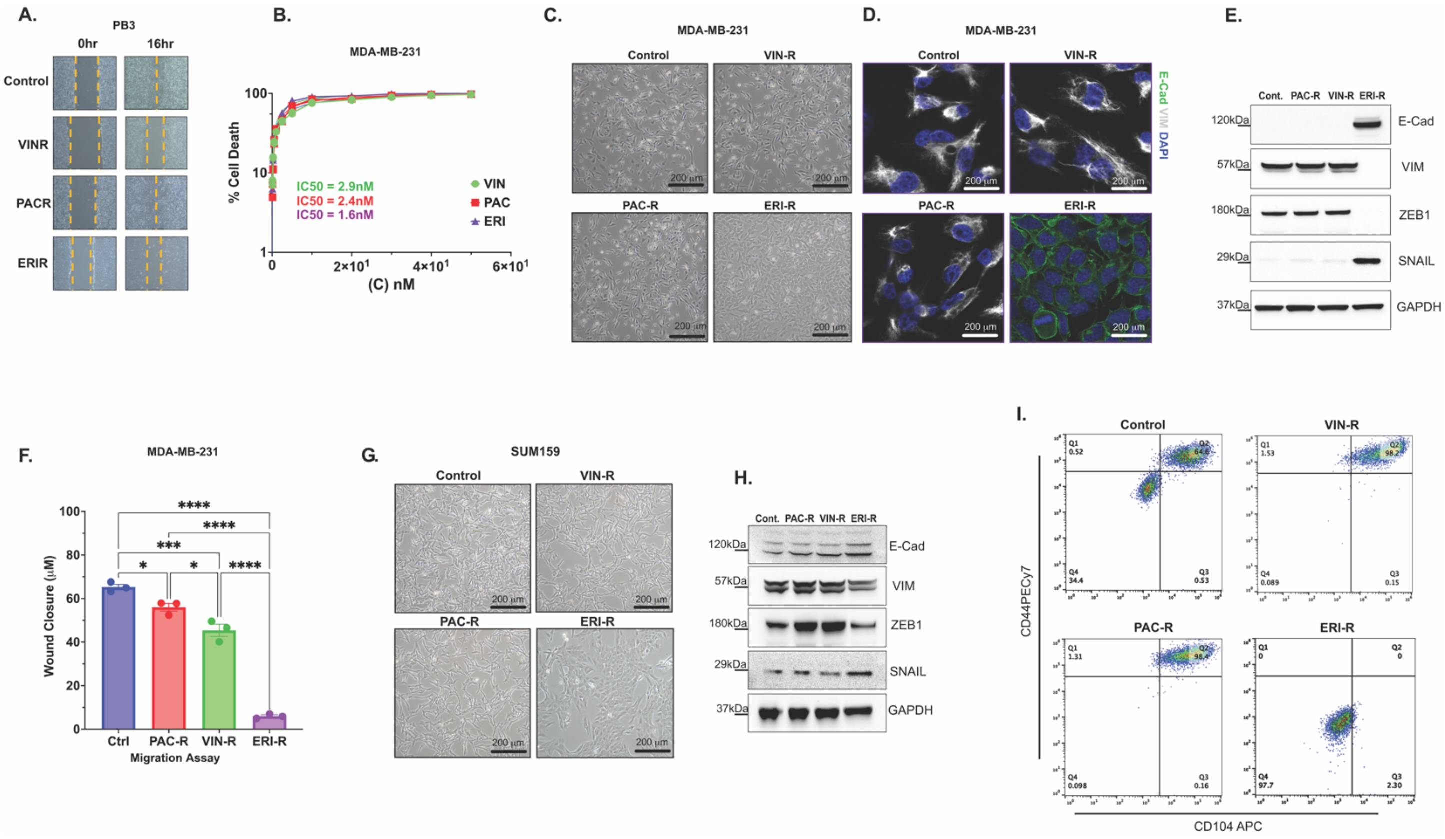


**Supplementary Figure 1: Eribulin induces MET in breast cancer cells.** (A) wound healing scratch assay was carried out in PB3 cells to assess the invasive properties of cells over 16 hours. (B) Dose response curves in MDA-MB-231 cells to calculate $IC_{50}$ values for eribulin, paclitaxel, and vinorelbine treatment. Estimation of EMT state was carried out by (C) morphological assessment of brightfield images, (D) immunofluorescence of canonical markers E-cadherin (green) and Vimentin (white), and (E) immunoblotting to estimate protein levels of EMT markers in the MDA-MB-231 parental line and resistant counterparts. (F) wound healing scratch assay was carried out in MDA-MB-231 cells to assess the invasive properties of cells over 16 hours. EMT state was estimated by (G) morphological assessment of brightfield images and (H) immunoblotting to estimate protein levels of EMT markers in the SUM159PT parental line and resistant counterparts. (I) Flow cytometry-based quantification of CD104 and CD44 expression in MDA-MB-231 parental cells and resistant clones.

#### Supplementary Fig 2

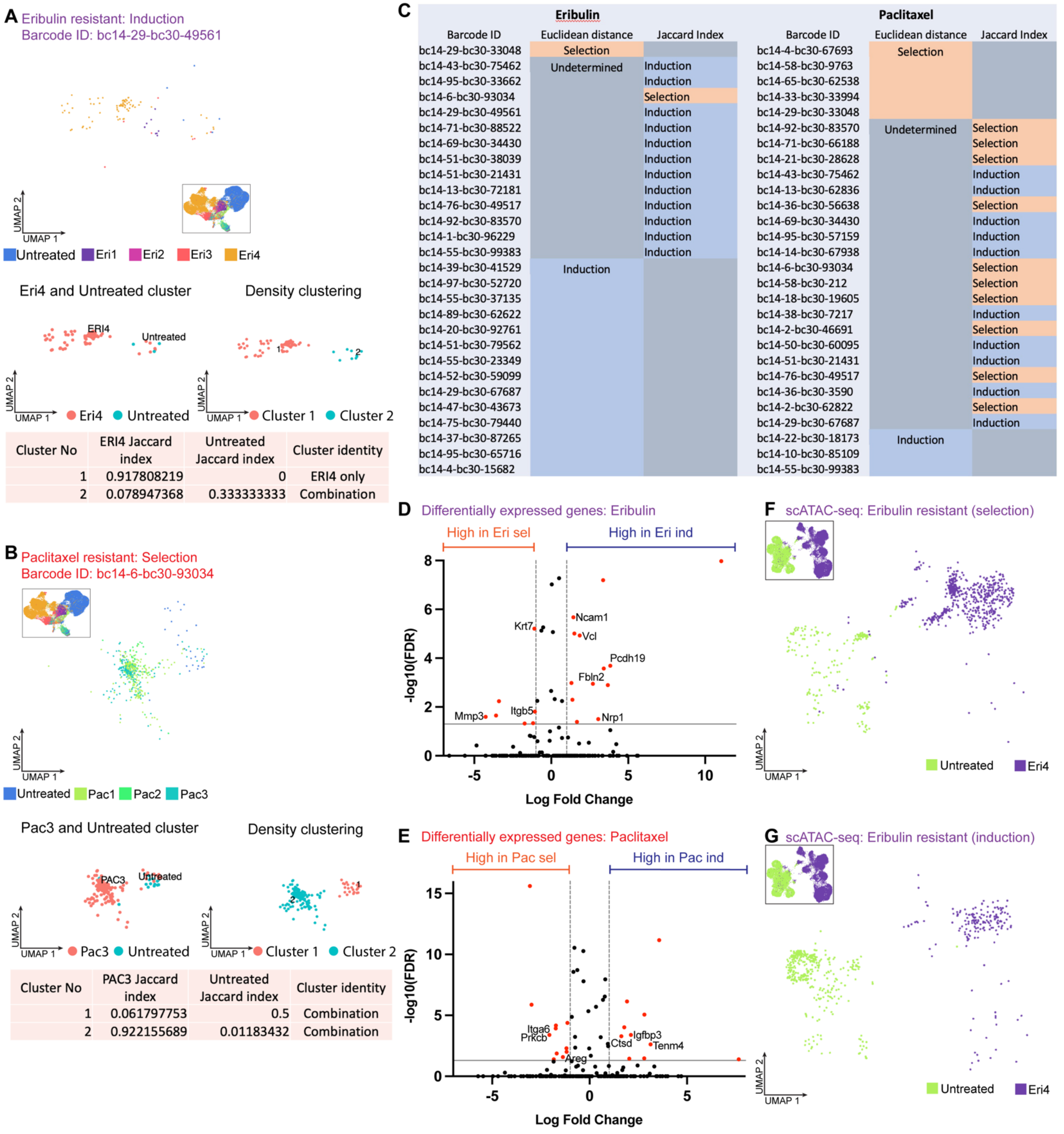


A:

| Cluster No | ERI4 Jaccard index | Untreated Jaccard index | Cluster identity |
|---|---|---|---|
| 1 | 0.917808219 | 0 | ERI4 only |
| 2 | 0.078947368 | 0.333333333 | Combination |

B:

| Cluster No | PAC3 Jaccard index | Untreated Jaccard index | Cluster identity |
|---|---|---|---|
| 1 | 0.061797753 | 0.5 | Combination |
| 2 | 0.922155689 | 0.01183432 | Combination |

C:

| Eribulin | | | Paclitaxel | | |
|---|---|---|---|---|---|
| Barcode ID | Euclidean distance | Jaccard Index | Barcode ID | Euclidean distance | Jaccard Index |
| bc14-29-bc30-33048 | Selection | | bc14-4-bc30-67693 | Selection | |
| bc14-43-bc30-75462 | Undetermined | Induction | bc14-58-bc30-9763 | | |
| bc14-95-bc30-33662 | | Induction | bc14-65-bc30-62538 | | |
| bc14-6-bc30-93034 | | Selection | bc14-33-bc30-33994 | | |
| bc14-29-bc30-49561 | | Induction | bc14-29-bc30-33048 | | |
| bc14-71-bc30-88522 | | Induction | bc14-92-bc30-83570 | Undetermined | Selection |
| bc14-69-bc30-34430 | | Induction | bc14-71-bc30-66188 | | Selection |
| bc14-51-bc30-38039 | | Induction | bc14-21-bc30-28628 | | Selection |
| bc14-51-bc30-21431 | | Induction | bc14-43-bc30-75462 | | Induction |
| bc14-13-bc30-72181 | | Induction | bc14-13-bc30-62836 | | Induction |
| bc14-76-bc30-49517 | | Induction | bc14-36-bc30-56638 | | Selection |
| bc14-92-bc30-83570 | | Induction | bc14-69-bc30-34430 | | Induction |
| bc14-1-bc30-96229 | | Induction | bc14-95-bc30-57159 | | Induction |
| bc14-55-bc30-99383 | | Induction | bc14-14-bc30-67938 | | Induction |
| bc14-39-bc30-41529 | Induction | | bc14-6-bc30-93034 | | Selection |
| bc14-97-bc30-52720 | | | bc14-58-bc30-212 | | Selection |
| bc14-55-bc30-37135 | | | bc14-18-bc30-19605 | | Selection |
| bc14-89-bc30-62622 | | | bc14-38-bc30-7217 | | Induction |
| bc14-20-bc30-92761 | | | bc14-2-bc30-46691 | | Selection |
| bc14-51-bc30-79562 | | | bc14-50-bc30-60095 | | Induction |
| bc14-55-bc30-23349 | | | bc14-51-bc30-21431 | | Induction |
| bc14-52-bc30-59099 | | | bc14-76-bc30-49517 | | Selection |
| bc14-29-bc30-67687 | | | bc14-36-bc30-3590 | | Induction |
| bc14-47-bc30-43673 | | | bc14-2-bc30-62822 | | Selection |
| bc14-75-bc30-79440 | | | bc14-29-bc30-67687 | | Induction |
| bc14-37-bc30-87265 | | | bc14-22-bc30-18173 | Induction | |
| bc14-95-bc30-65716 | | | bc14-10-bc30-85109 | | |
| bc14-4-bc30-15682 | | | bc14-55-bc30-99383 | | |

**Supplementary Figure 2: Clonal dynamics following drug treatment.** (A) Representative plots of how the determination of selection vs. induction was made based on Jaccard Index. (A) Representative plot of a combination of induction and selection within the same barcode. For this barcode, certain clusters are resistant by induction (no untreated cells in the final treatment cluster), and others are resistant by selection (at least one untreated cell in the final treatment cluster). The cluster with the largest number of resistant cells are used to determine the mode of resistance. In this example, cluster 1 has the largest number of Eri4 cells, and these cells are resistant by induction, therefore, this barcode was assigned as resistant by induction. (B) Representative plots of barcodes resistant by selection. Untreated cells and PAC3 with this barcode share the same clusters as observed by the Jaccard index score. (C) The levels of induction or selection were determined by breakdown of both Euclidean distance and Jaccard index calculations. Volcano plots showing differentially expressed genes that may predict induction or selection resistance upon (D) eribulin and (E) paclitaxel treatment. Gene expression profiles of untreated cells with the same barcode eribulin induction resistant cells and paclitaxel induction resistant cells (highest 10 Euclidean distance values) were compared with untreated cells with the same barcode as eribulin and paclitaxel selection resistant cells (lowest 10 Euclidean distance values). Significant genes were determined by an FDR of <0.05, and a fold change of >2. scATAC-seq UMAP projection of untreated (F) eribulin resistant cells by selection and (G) eribulin treated cells resistant by induction (highest 10 Euclidean distance values).

#### Supplementary Fig 3

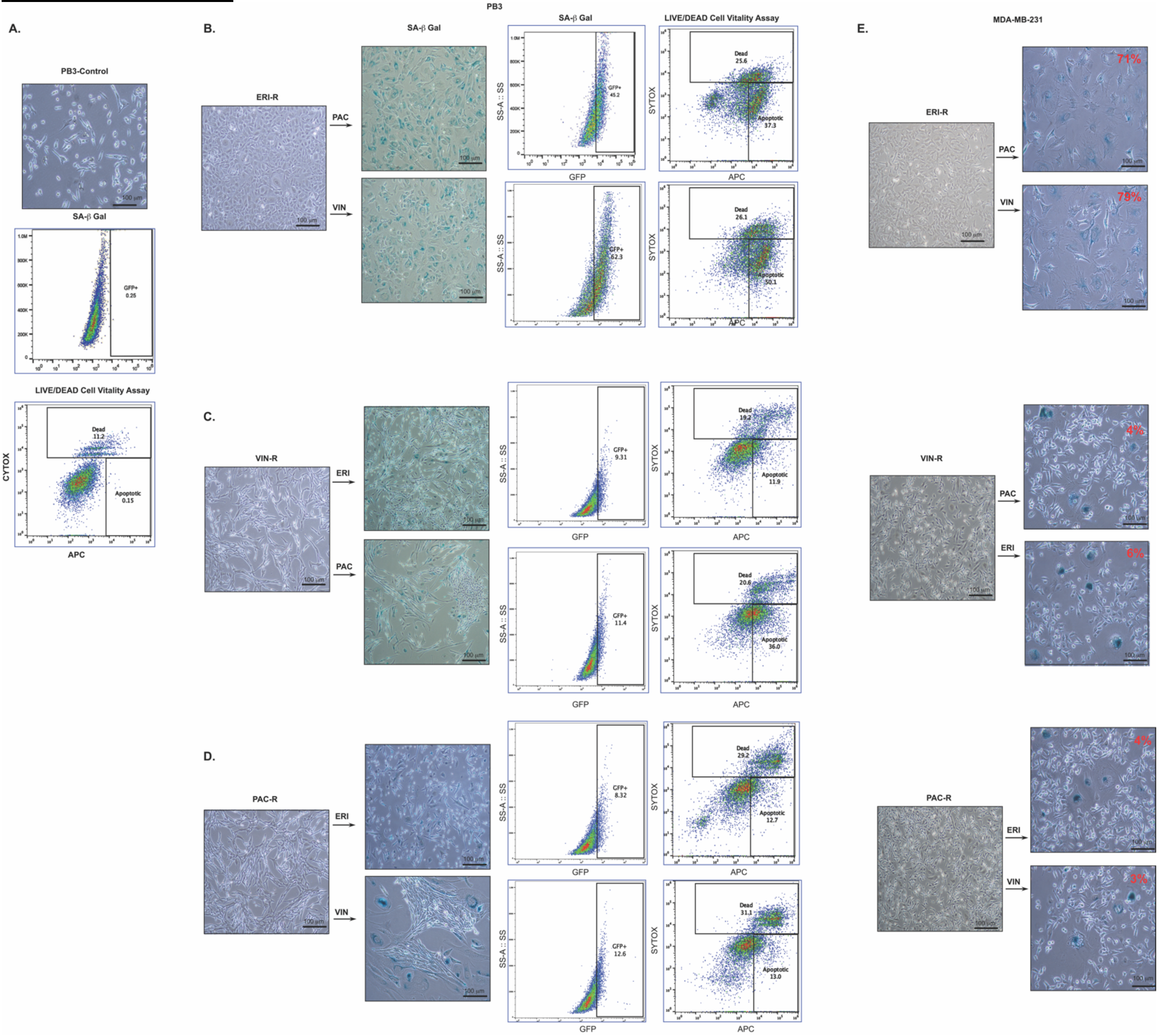


**Supplementary Figure 3: Eribulin pre-treatment induces sensitization to subsequent therapy.** (A) Quantification of live-dead and apoptotic cells using SYTOX in PB3 parental cells. Senescence-associated beta-galactosidase assay (brightfield images and flow cytometry panel on left) and quantification of live-dead and apoptotic cells using SYTOX (flow cytometry panel on right) to evaluate drug response of PB3-derived (B) ERI-R cells to paclitaxel and vinorelbine treatment, (C) PAC-R cells to eribulin and vinorelbine treatment, and (D) VIN-R cells to paclitaxel and eribulin treatment. (E) Senescence-associated beta-galactosidase assay to determine drug response of MDA-MB-231 derived ERI-R cells to paclitaxel and vinorelbine treatment, PAC-R cells to eribulin and vinorelbine treatment, and VIN-R cells to paclitaxel and eribulin treatment.

#### Supplementary Fig 4

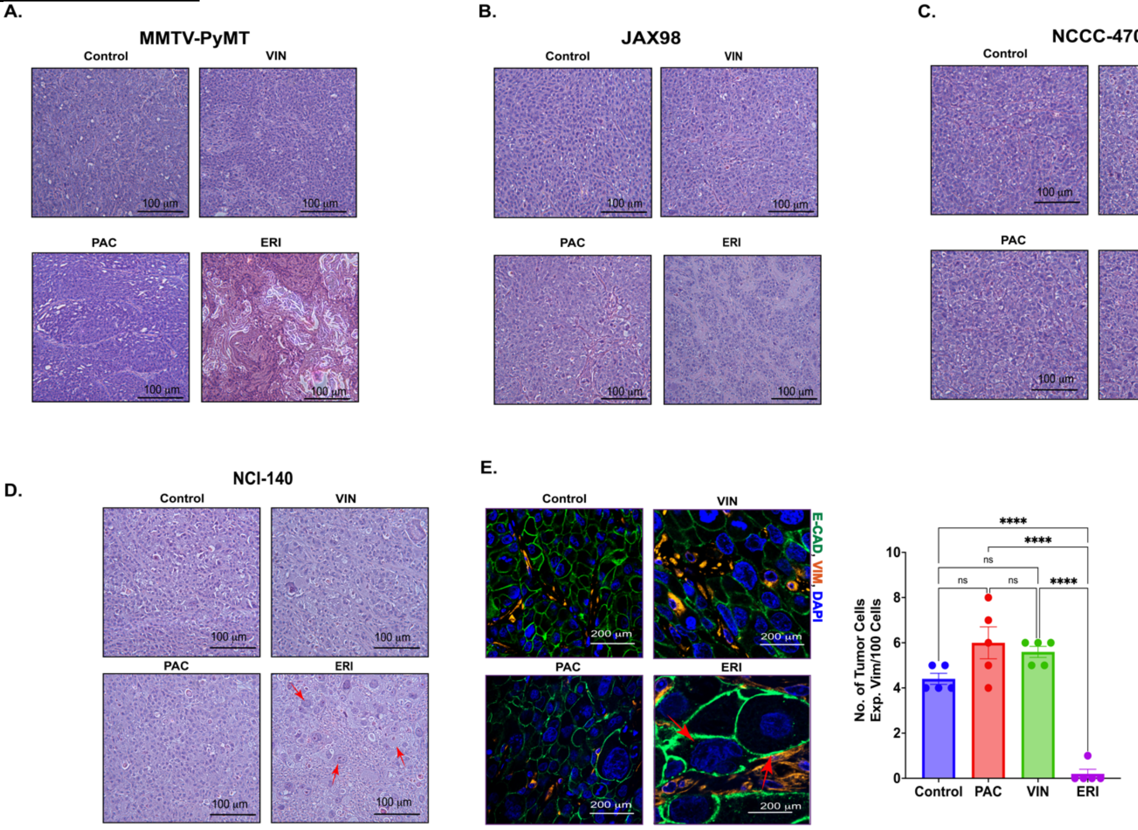


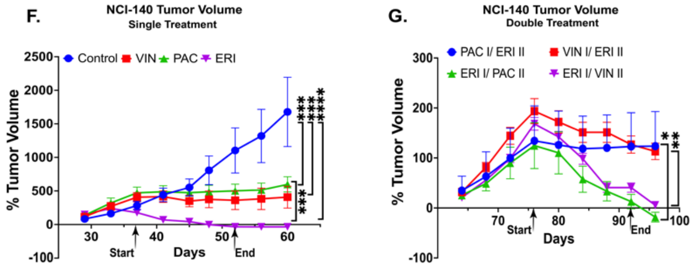


H.

| NCI-140 Metastasis | | | | |
|---|---|---|---|---|
| Group | Lung | Liver | Ovary | lymph node |
| Control | 1/5 | 0/5 | 5/5 | 1/5 |
| E1P2 | 0/5 | 0/5 | 0/5 | 0/5 |
| E1V2 | 0/5 | 0/5 | 0/5 | 0/5 |
| P1E2 | 0/5 | 0/5 | 1/5 | 0/5 |
| V1E2 | 0/5 | 0/5 | 1/5 | 0/5 |

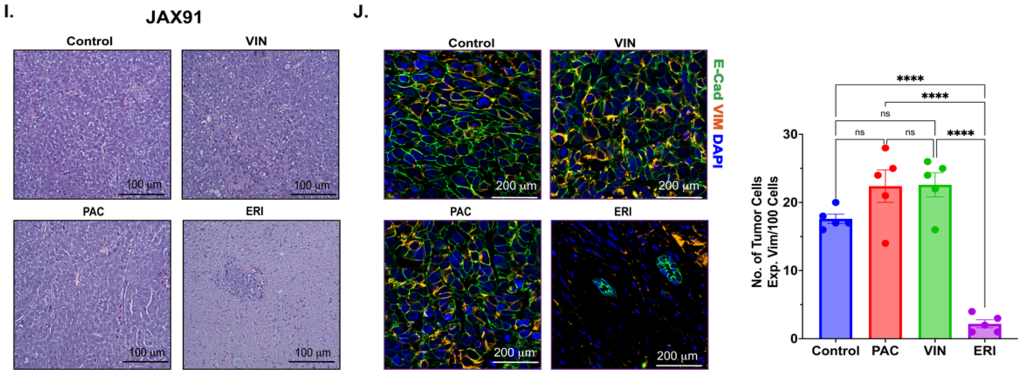


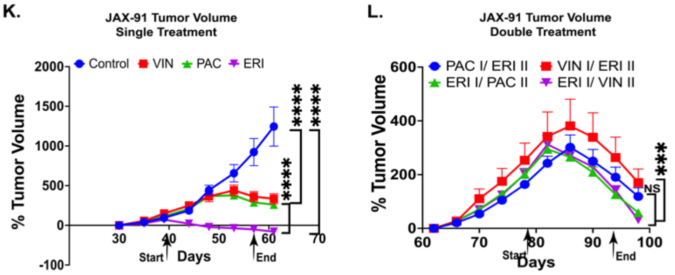


M.

| JAX-91 | | | | |
|---|---|---|---|---|
| Group | Lung | Liver | Ovary | lymph node |
| Control | 0/0 | 3/5 | 4/5 | 4/5 |
| E1P2 | 0/0 | 0/0 | 1/5 | 2/5 |
| E1V2 | 0/0 | 0/0 | 0/0 | 1/5 |
| P1E2 | 0/0 | 0/0 | 4/5 | 4/5 |
| V1E2 | 0/0 | 0/0 | 2/5 | 2/5 |

**Supplementary Figure 4: MET induction is accompanied by robust tumor regression and reduced metastatic burden.** H&E histology representative of (A) MMTV-PyMT tumor-bearing mice, (B) JAX-98, and (C) NCCC-470 were treated with vinorelbine, paclitaxel, and eribulin. Triple negative breast cancer derived PDXs, NCI-140 (naïve) and JAX-91 (pretreated) were treated with vinorelbine, paclitaxel, and eribulin to assess (D and I) H&E histology examination, (E and J) EMT status using immunohistofluorescence of canonical EMT markers E-cadherin (green) and Vimentin (red). (F and K) Tumor growth rate of single drug regimens, (G and L) and sequential double treatment as well as (H and M) distribution patterns of tumor metastasis from the sequential double treatment to the lung, respectively. ($*p <0.05$, $**p < 0.001$, $***p > 0.0005$, $****p < 0.0001$, Two-Way ANOVA, n=5).

**Supplementary Fig 5**

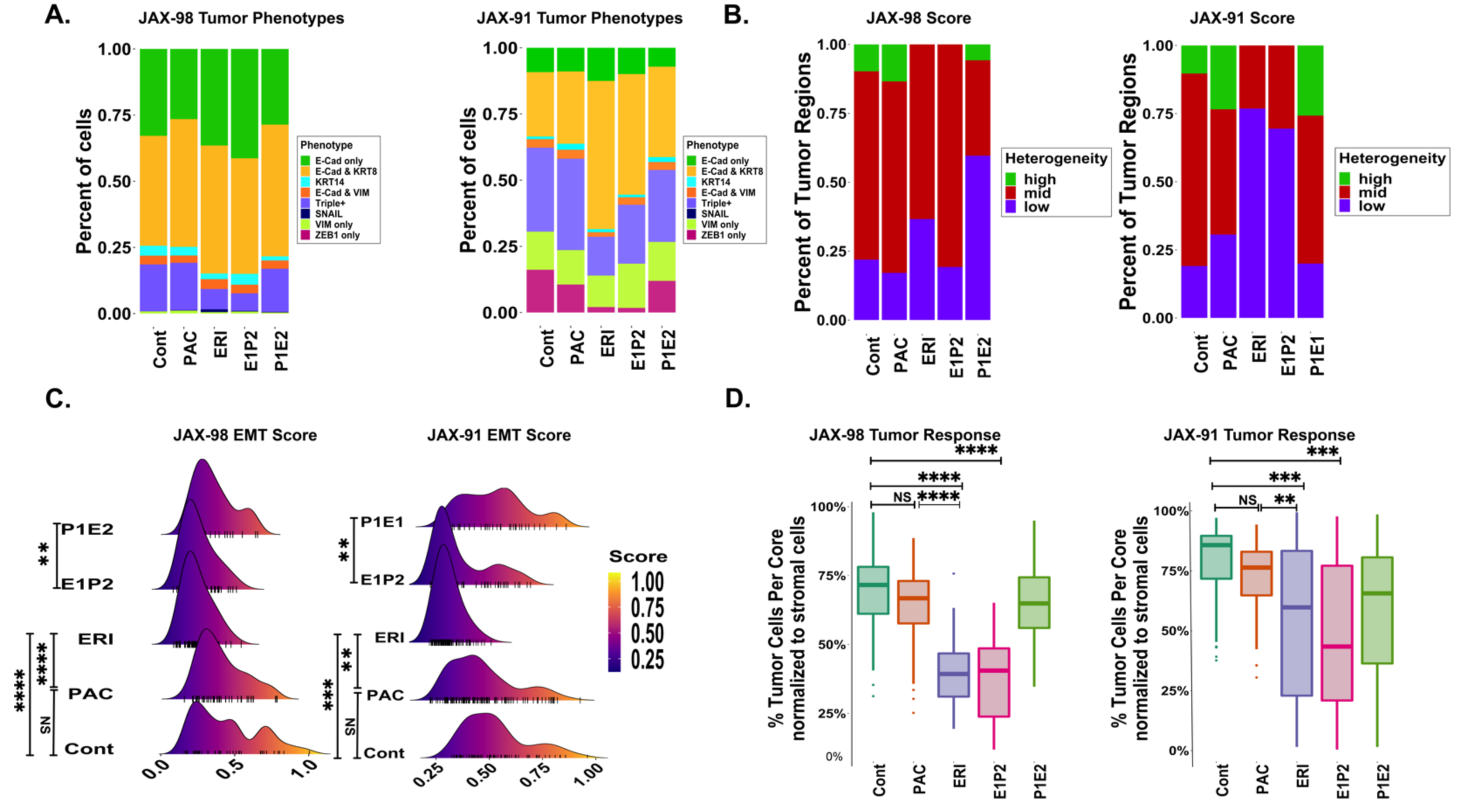


**Supplementary Figure 5: MET induction leads to the reduction in epithelial-mesenchymal heterogeneity of tumors.** Multiplexed, multi-round TSA staining of EMT markers in triple negative breast cancer derived PDXs (JAX-98 and JAX-91) treated with paclitaxel and eribulin as a single and sequential double treatment to evaluate (A) distribution of tumor phenotypes by calculating the percent of tumor cells expressing different EMT markers, (B) entropy-based tumor heterogeneity levels, (C) EMT score, and (D) Tumor response to drug treatment. (∗p <0.05, ∗∗p < 0.001, ∗∗∗p > 0.0005, ∗∗∗∗p < 0.0001, Two-Way ANOVA, n=5).

#### Supplementary Fig 6

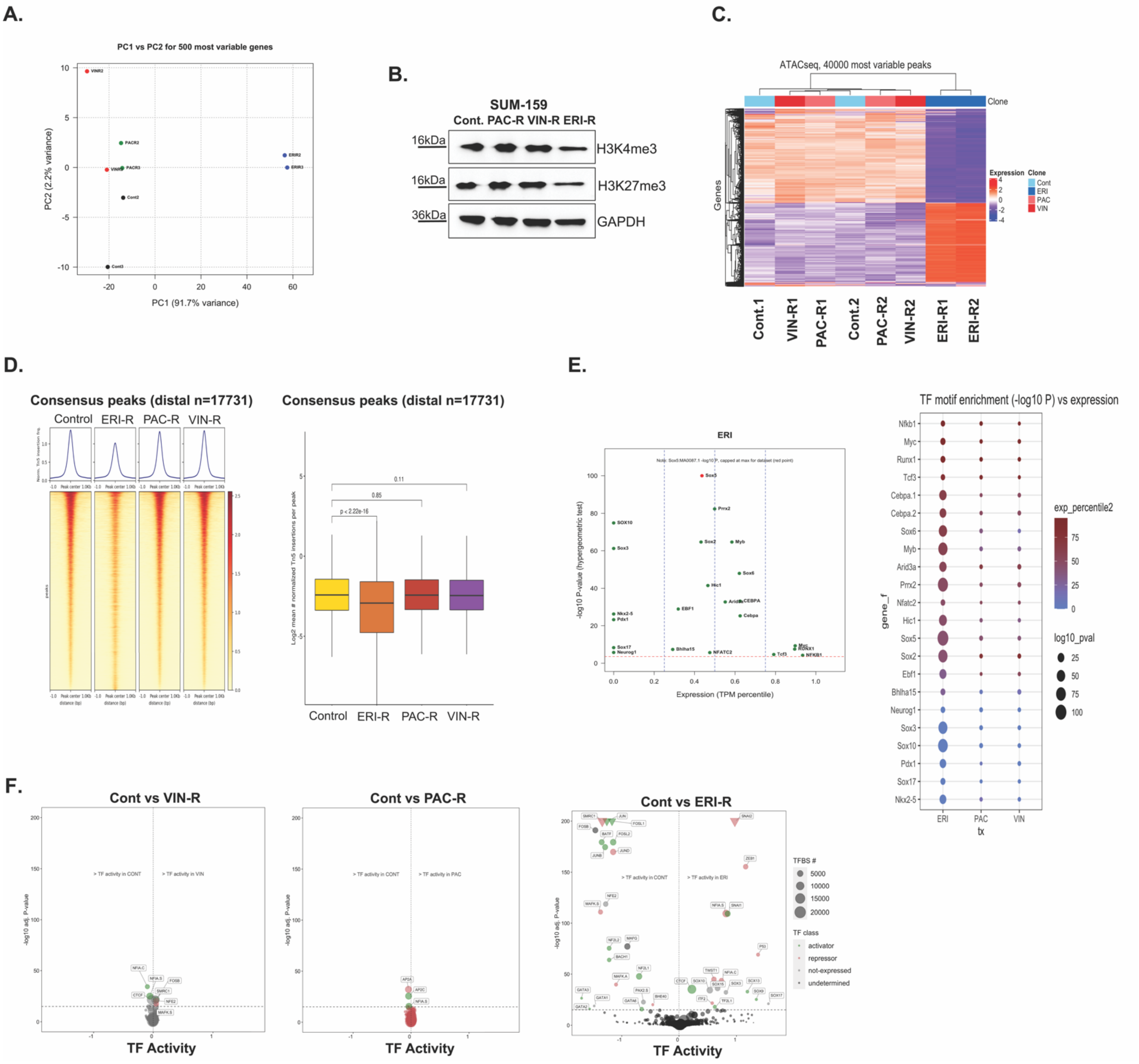


**Supplementary Figure 6: Treatment with eribulin induces a shift in the chromatin profile of cancer cells.** (A) Principal Component Analysis (PCA) of the 500 most variable genes between PB3 parental and resistant clones from RNA-seq. (B) Immunoblotting showing levels of H3K4me3 and H3K27me3 evaluated in SUM159PT parental and resistant clones. ATAC-sequencing was performed in PB3 parental and resistant clones to explore chromatin accessibility at different regions (C) heat map of most variable peaks and, (D) peak accessibility surrounding all consensus distal-associated regions. (E) TF motifs enrichment evaluated in PB3 ERI-R cells, the left panel shows highly significant active and inactive transcription factors. The X-axis shows TF targets expression, and the y-axis shows -Log10 p-values of each transcription factor (left panel). The right panel shows TF motif enrichment over target expressions circle size indicates the p-value, and the circle color shows the target genes' percentile expression across indicated samples. (F) Advanced volcano plot of active and inactive transcription factors determined by diffTF from ATAC-seq. Transcription factor classification, displayed in bubble color, and the number of transcription factor binding sites used to determine TF activity plotted as bubble size. TFs in triangles are highly significant.

#### Supplementary Fig 7

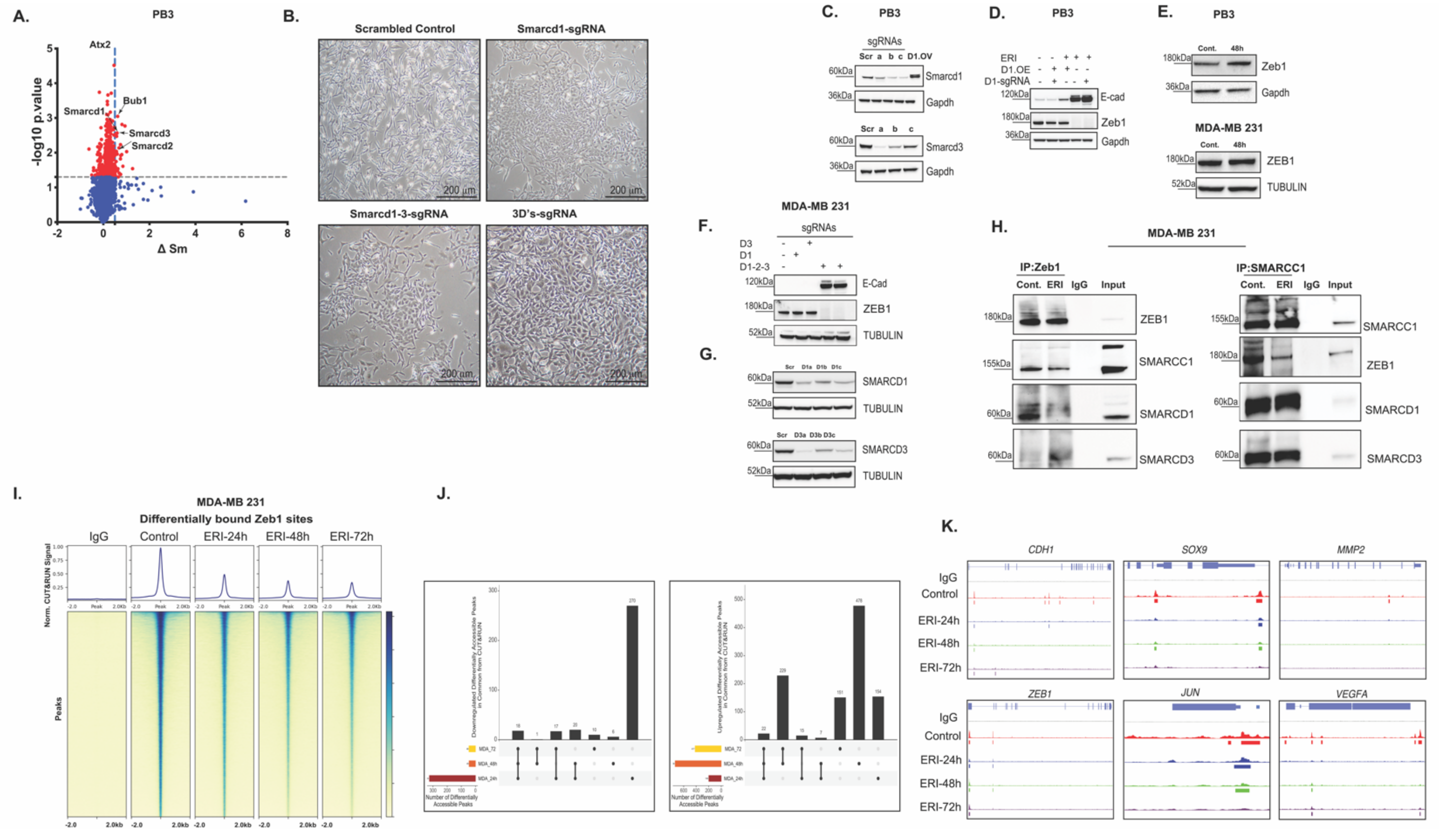


**Supplementary Figure 7: ZEB1-SWI/SNF interactions are necessary for maintenance of a mesenchymal state**. (A) Proteome Integral Solubility Alteration of PB3 cells treated with 600nM eribulin for 4 hours to explore the non-tubulin targets of eribulin. Indicated proteins show the highest delta Sm and p-value. (Delta Sm > 0.5 and $p < 0.05$, T-test), and estimation of EMT state as carried out by (B) morphological assessment of brightfield images. (C) CRISPR/Cas9-mediated deletion of Smarcd1, and Smarcd3 genes to explore the knockdown efficiency in PB3 cells. Three different sgRNAs per gene (a, b, and c) were used. (D) SMARCD1 overexpressed in PB3, PB3-ERI-R control and their Smarcd1.kd counterparts to explore the EMT state using canonical EMT makes E-cad and Zeb1. (E) Immunoblot analysis of PB3 (top) and MDA-MB-231 (bottom) to explore the expression levels of Zeb1/ZEB1 48 hours after eribulin treatment. (F) EMT state was evaluated using immune blot analysis in MDA-MB-231 cells with indicated conditions. (G) Western blot analysis explores the knockdown efficiency of SMARCD1 and SMARCD3 cells in MDA-MB-231 cells. Three different sgRNAs per gene (a, b, and c) were used. (H) Co-immunoprecipitation using an anti-ZEB1 antibody (left panel) and anti-SMARCC1 antibody and probed for indicated proteins (right panel) in MDA-MB-231 parental and eribulin treated cells. The genome-wide occupancy of ZEB1 binding sites in MDA-MB-231 parental cells and those treated with eribulin for 24, 48, and 72 hours was evaluated using CUT&RUN. (I) The average CUT&RUN enrichment profile (top) and heatmap illustrating the CUT&RUN signal 2 kb up-and downstream of the different peaks, (J) UpSet plot of differentially accessible peaks at the indicated time points and (K) signal track of Zeb1 target genes accessibly.
